# Transcription initiation defines kinetoplast RNA boundaries

**DOI:** 10.1101/350256

**Authors:** François M. Sement, Takuma Suematsu, Liye Zhang, Tian Yu, Lan Huang, Inna Aphasizheva, Ruslan Aphasizhev

## Abstract

Mitochondrial genomes are often transcribed into polycistronic primary RNAs punctuated by tRNAs whose excision defines mature RNA boundaries. Although kinetoplast DNA lacks tRNA genes, it is commonly held that monophosphorylated 5′-ends of functional molecules typify precursor partitioning by an unknown endonuclease. To the contrary, we demonstrate that in *Trypanosoma brucei* individual mRNAs and rRNAs are independently synthesized as 3′ extended precursors. The transcription-defined 5′ terminus is converted into monophosphorylated state by the 5′ pyrophosphohydrolase complex, termed PPsome, which is activated by RNA editing substrate binding complex (RESC). Most guide RNAs lack PPsome recognition sites and, therefore, remain triphosphorylated. We provide evidence that both 5′ pyrophosphate removal and 3′ adenylation are essential for mRNA stabilization. Furthermore, we uncover a mechanism by which antisense RNA-controlled 3′-5′ exonucleolytic trimming defines mRNA 3′-end. We conclude that mitochondrial mRNAs and rRNAs are transcribed and processed as insulated units irrespective of their genomic location.

**Significance:** It is commonly held that in trypanosomes both mitochondrial DNA strands are transcribed into polycistronic precursors. These primary RNAs are presumably partitioned into individual pre-mRNAs by a “cryptic” endonuclease. We challenged the polycistronic transcription/ endonuclease model after revealing precursor processing by 3′-5′ degradation. This work demonstrates individual transcription of each gene and mRNA 5′-end definition by the first incorporated nucleotide triphosphate. We have uncovered the stabilizing role of 5′ triphosphate to monophosphate conversion and identified a protein complex responsible for this reaction. We have discovered antisense noncoding RNA originating near mRNA 3′ end and showed that a duplex formation modulates exonuclease activity to delimit the mature 3′ end. Collectively, our findings reveal mechanisms by which transcription defines both mRNA termini.

## Introduction

Notwithstanding their monophyletic origin, present-day mitochondria display an inexplicable diversity of transcriptional, RNA processing, and translation mechanisms. In animals and fungi, mitochondrial DNA is transcribed into polycistronic primary RNAs which are cleaved internally (1, 2), while in plants diverse *cis*-elements recruit RNA polymerases to individual genes (3). As suggested by the “tRNA punctuation” model, pre-mRNAs are liberated from polycistronic precursors via excision of flanking tRNAs by RNases P and Z (4, 5). The causative agent of African sleeping sickness, *Trypanosoma brucei*, maintains a bipartite mitochondrial genome composed of a few ~23-kb maxicircles and thousands of ~1-kb minicircles. Maxicircles encode 9S and 12S rRNAs, 18 tightly packed protein genes, and a single guide RNA; the minicircles produce gRNAs required for U-insertion/deletion mRNA editing (6). Interestingly, mature rRNA and mRNA 5′ termini are monophosphorylated (5′P) while gRNAs retain triphosphate (5′3P) characteristic of the transcription start site (7). A putative maxicircle transcription initiation region has been mapped to the major strand ~1200 nucleotides upstream of the 12S rRNA, and the transcription is believed to proceed polycistronically (8, 9). Although the absence of mitochondrial tRNA genes (10) negates the “tRNA punctuation” scenario, it is commonly held that an unknown endonuclease cuts between functional sequences (11, 12). However, studies of multiple endonucleases have neither provided evidence for the “cryptic” processing activity, nor identified demarcation elements positioned between coding sequences (13–15).

In contrast to maxicircle transcripts, guide RNAs are synthesized from dedicated promoters as ~1kb precursors and processed by 3′-5′ exonucleolytic trimming (16, 17). This reaction is carried out by DSS1 3′-5′ exonuclease (18) acting as subunit of the mitochondrial 3′ processome (MPsome) (16). Recently, we established that rRNA and mRNA precursors accumulate upon knockdown of the MPsome’s components (19). Hence, 3′-5′ trimming appears to be the major, if not the only, pathway for nucleolytic processing of primary RNAs. This modus operandi, however, would be incongruent with a polycistronic precursor containing several coding sequences: Only the 5′ region can be converted into pre-mRNA that is competent for polyadenylation, editing and translation. Furthermore, any conceivable mechanism ought to account for the monophosphorylated 5′ termini and homogenous 3′-ends.

In this work, we demonstrate that mRNA and rRNA 5′-ends are defined by transcription initiation while the 3′-extended primary transcripts encroach into downstream genes. The 5′3P moiety is converted into 5′P by MERS1 NUDIX (<u>nu</u>cleoside <u>di</u>phosphates linked to <u>x</u> (any moiety)) hydrolase (20). Along with a MERS2 pentatricopeptide repeat factor and MERS3 subunit lacking any motifs, MERS1 constitutes a 5′ pyrophosphohydrolase complex, termed the PPsome. Catalytically inactive as individual protein, the PPsome-imbedded MERS1 displays a hydrolase activity, which specifically targets mRNA 5′ termini and is further stimulated by the <u>R</u>NA <u>e</u>diting <u>s</u>ubstrate binding <u>c</u>omplex (RESC, (21)). The PPsome apparently functions as a “protein cap” to stabilize monophosphorylated mRNAs by interacting with the polyadenylation complex to tether 5′ and 3′ termini. Finally, we determine the mechanism by which the antisense non-coding RNAs modulate 3′-5′ degradation thereby defining the 3′-ends of maxicircle-encoded mRNAs.

## Results

### Kinetoplast Genes Are Transcribed as Independent Units

To investigate mitochondrial RNA polymerase (MTRNAP) occupancy of the maxicircle DNA, we developed the <u>k</u>inetoplast <u>a</u>ffinity <u>p</u>urification – sequencing (KAP-Seq) protocol. The C-terminally TAP-tagged MTRNAP was expressed in an insect (procyclic) developmental form of the parasite and verified to have been targeted to the mitochondrial matrix without appreciably impacting cell growth (Fig. S1A, B). The 170 kDa MTRNAP-TAP was incorporated into an ~900 kDa (22S) complex (Fig. 1A), which was resistant to RNase treatment (Fig. S1C). Live cells were crosslinked with formaldehyde (Fig. 1B) and DNA was fragmented by focused sonication (Fig. S1D). At the ~100 bp resolution achieved with KAP-Seq (Fig. 1C), the MTRNAP binding was detected predominantly within conserved gene-containing region (Fig. 1D). This trend is particularly instructive for adjacent genes, such as 9S and 12S mt-rRNAs or the *ND7-CO3-cyb-A6* segment. Here, polycistronic transcription would be expected to correlate with a uniform MTRNAP progression. However, the *cyb* gene clearly shows a decreased amount of MTRNAP KAP-Seq reads compared to neighboring ND7, *CO3* and *A6*. In the segment devoid of annotated genes, a major occupancy peak matched the position of a previously mapped cryptic precursor originating ~1200 nt upstream of the 12S rRNA (8). Hence, the MTRNAP occupancy appears to correlate with positioning of individual rRNA and protein genes, and transcripts of unknown function.

**Fig. 1.**
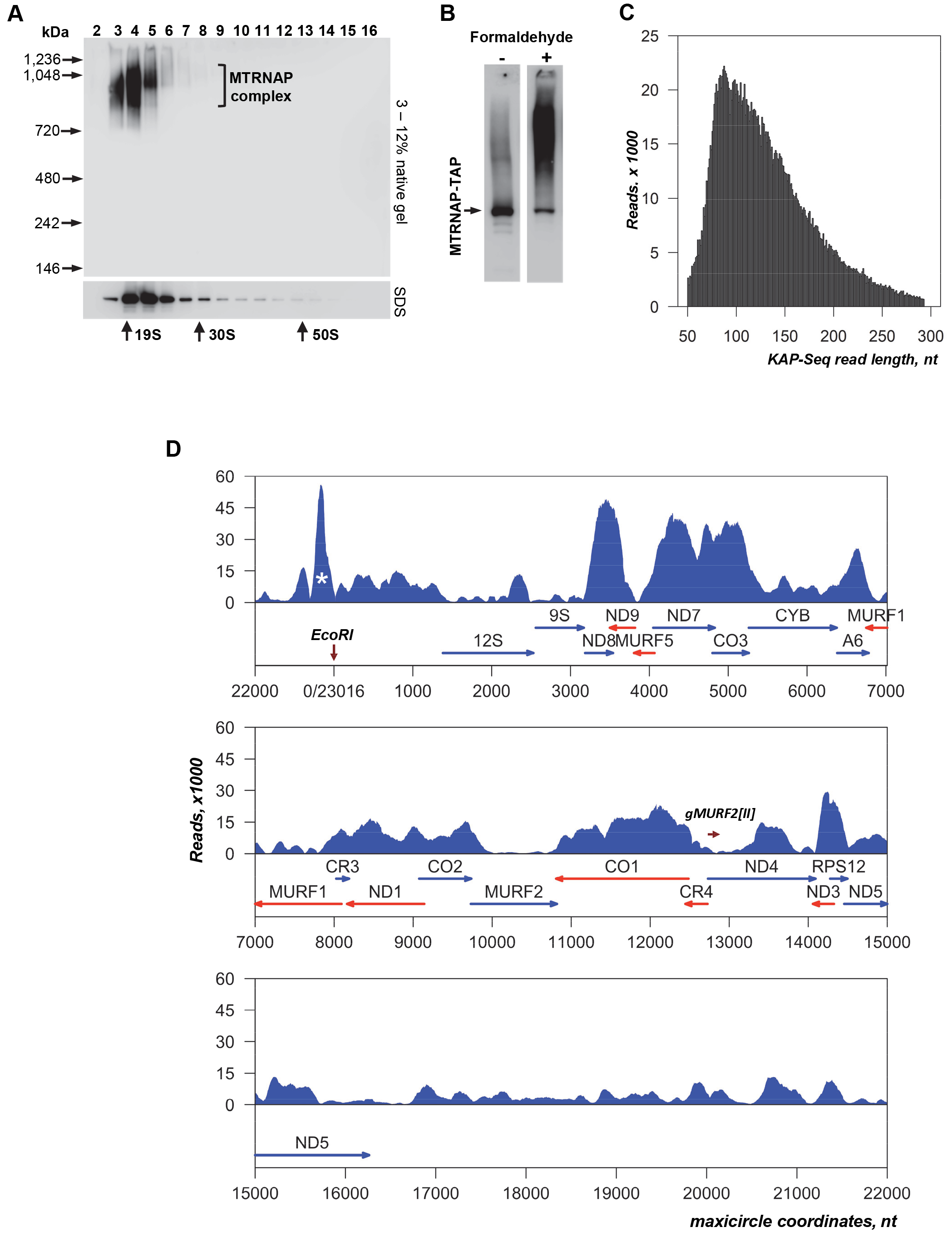
RNA polymerase occupancy conforms to individual gene boundaries. (A) Glycerol gradient fractionation of MTRNAP complex. Lysate from cells expressing TAP-tagged MTRNAP was separated in 10% – 30% glycerol gradient. Each fraction was resolved on native and denaturing gels. TAP fusion protein was detected by immunoblotting. Positions of native molecular mass and sedimentation markers are indicated. (B) Formaldehyde-induced crosslinking of MTRNAP. Live cells expressing MTRNAP-TAP were crosslinked with formaldehyde, extracted and separated on SDS gel. (C) Length distribution of KAP-Seq reads that mapped to the maxicircle. (D) MTRNAP occupancy of the maxicircle. Genes located on major and minor strands are diagrammed by blue and red arrows, respectively. A single maxicircle-encoded guide RNA, gMURF2[II], lies within, but is transcribed independently from ND4 mRNA (42). Read counts and maxicircle coordinates (GenBank: M94286.1) are shown starting at the unique EcoR I site. The peak in the non-coding region upstream of 12S rRNA corresponding to the major cryptic precursor identified by (8) is marked by asterisk.

### Transcription Initiation Defines the 5′ End

To corroborate independent transcription of individual genes, we performed *in vivo* UV-crosslinking affinity purification – sequencing (CLAP-Seq) to identify nascent RNAs bound to transcribing RNA polymerase (Fig. 2A). In this application, cell extract was treated with RNases A and T1 to fragment RNA while isolated RNA-protein adducts were digested on beads with 3′-5′ exonuclease RNase R. The latter step reduces overall background by degrading RNAs with 3′ ends unprotected by MTRNAP. In agreement with the DNA binding profile, the CLAP-Seq reads distribution along the maxicircle coding region largely concurred with gene boundaries and often showed accumulation at 5′ and 3′ ends of individual genes (Fig. 2B). Remarkably, omitting RNase A/T1 digestion enriched 5′ regions among MTRNAP-bound mRNAs (Fig. 2C). Statistical analysis demonstrated a significant increase in number of reads covering the 5′ termini of 18 out of 20 annotated maxicircle transcripts (Fig. 2D and Dataset S1, χ^2^ test).

**Fig. 2.**
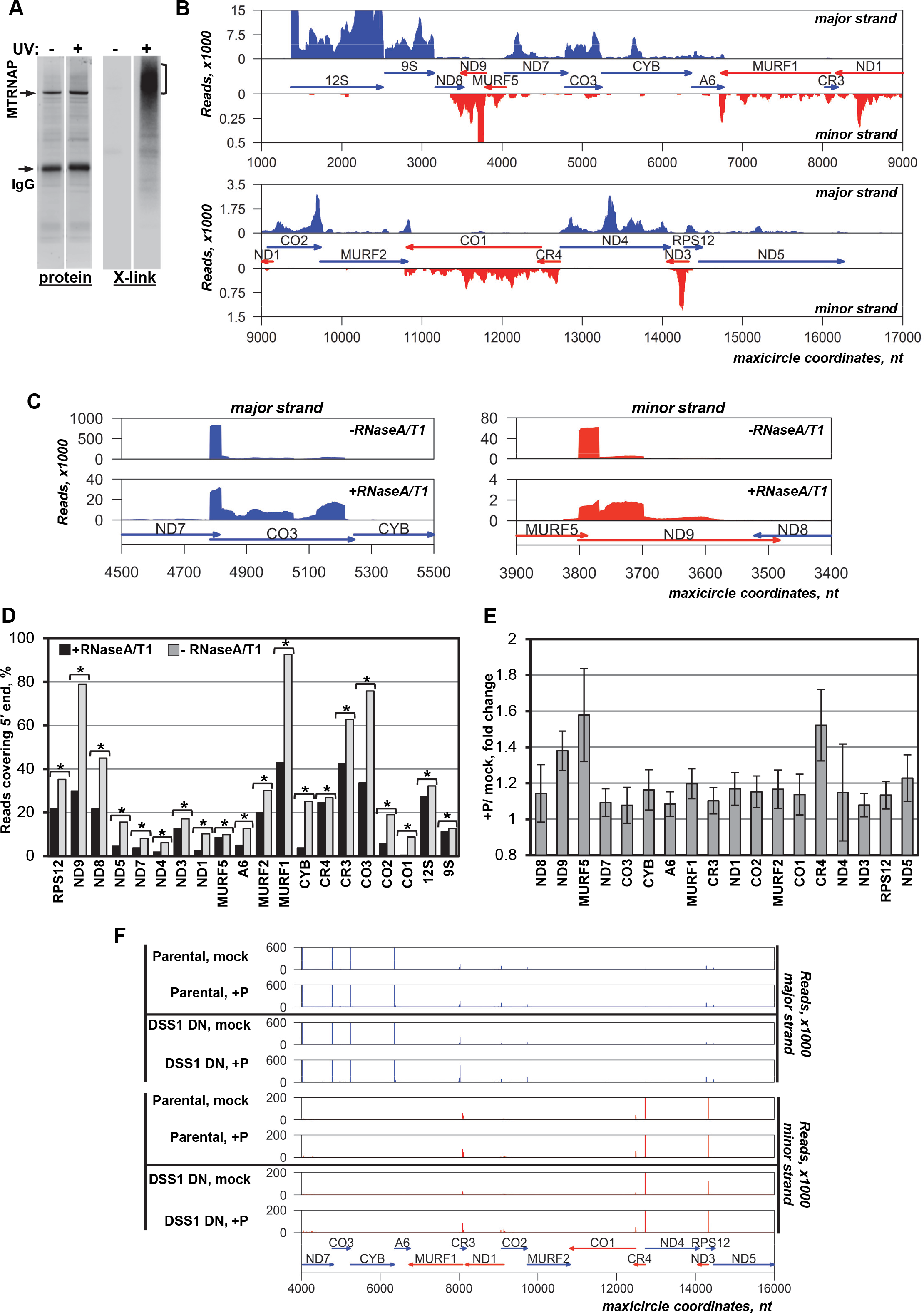
Mature 5′ termini correspond to transcription initiation sites. (A) MTRNAP-RNA UV-crosslinking affinity purification (CLAP). The MTRNAP-TAP purification from mock-treated (−) or UV-irradiated (+) parasites was accompanied by RNase A and T1 fragmentation in extract, treatment with RNase R on beads, and radiolabeling of the crosslinked RNA. Adducts were separated on SDS PAGE, transferred onto nitrocellulose membrane, and visualized by Sypro Ruby staining (protein) or by exposure to a phosphor storage screen (X-link). RNA was eluted from (−) and (+) areas indicated by bracket and sequenced. (B) Positioning of nascent RNAs. MTRNAP CLAP reads were mapped to the coding region of the maxicircle. Predicted genes on major (blue) and minor (red) strands, read counts and maxicircle coordinates are indicated. (C) Representative examples of reads re-distribution upon omitting RNase A/T1 treatment during MTRNAP CLAP. (D) Enrichment of mRNA and rRNA 5′ regions in MTRNAP-bound RNAs. The percentage of CLAP reads covering the 5′ terminus was calculated for each gene. Asterisk denotes a significant 5′-end increase (χ^2^ test, p < 0.01). (E) Detection of individual triphosphorylated transcripts. The polyphosphatase-dependent 5′-end enrichment was calculated for parental cells as a ratio between normalized read counts in +P and mock-treated samples. Bars show standard deviations between four biological replicates. (F) Mapping the 5′-ends of maxicircle transcripts. The 5′-end of each RACE-Seq read was plotted on major (blue) and minor (red) maxicircle strands. Pearson’s correlation coefficients between global 5′ RACE profiles in parental and DSS1 DN backgrounds from four biological replicates are provided in Dataset S1, P-value <0.001.

The monophosphorylated state of the mature mRNA 5′-end has been exposed by molecular cloning (22). To directly test whether positions of primary and processed 5′ ends coincide, an RNA adapter was ligated to mock-and polyphosphatase-treated (+P) RNA. This reaction converts the 5′3P and 5′2P termini into 5′P substrate for T4 RNA ligase; therefore, an increase in read counts would reflect 5′3P and 5′2P occurrence. The gene-specific 5′ RACE-Seq libraries were constructed from the parental and the cell line conditionally expressing a dominant negative variant of DSS1 3′-5′ exonuclease (DSS1 DN). DSS1 repression causes accumulation of 3′ extended gRNA, rRNA, and mRNA precursors (16, 19). Hence, we reasoned that the positions of gene-specific transcription initiation sites should not change. In the parental cell line, the polyphosphatase-dependent gains in 5′ reads indicated that 10% – 45% of mRNA species retain the transcription-incorporated 5′ nucleoside triphosphate (Fig. 2E). Considering all mRNAs as a group, the combined gain in supporting reads is statistically significant with a P value of 0.006372 in a paired T-test. By examining 5′-ends at a single nucleotide resolution, we found that the sequences remained unaltered in DSS1 DN cells, while some transcripts became enriched in polyphosphatase-treated RNA (+P). Importantly, correlation analysis confirmed that the 5′ termini derived from mono-or di/triphosphorylated RNAs were virtually identical in parental and DSS1 DN cell lines (Dataset S1). These data corroborate synthesis of 5′-defined RNAs and an absence of endonucleolytic or 5′-3′ exonucleolytic processing at the 5′ termini (Fig. 2F). Collectively, *in vivo* analysis of MTRNAP occupancy, sequencing of nascent transcripts, determination of 5′-end positions and phosphorylation states demonstrate that individual genes are transcribed as independent units.

### Identification of the 5′ PPsome

Eukaryotic mRNAs are typically protected by a 5′ cap and 3′ poly(A) tail, while the 5′ triphosphate and 3′ stem-loop entities are critical for bacterial mRNA stability (23). In mitochondria of trypanosomes, a monophosphorylated 5′-end is apparently produced by PPi removal, but the cognate activity and functional implications of 5′3P into 5′P conversion are uncertain. To addresses these questions, we focused on NUDIX hydrolases that cleave nucleoside diphosphates linked to any moiety, including RNA. A survey of the *T. brucei* genome identified five potential NUDIX-like proteins (24), of which MERS1 is targeted to the mitochondrion. MERS1 was initially identified by co-purification with MRP1/2 RNA chaperones, but its function remained unclear (20). To place this enzyme into a functional context, we assessed MERS1 interactions by separating mitochondrial complexes on glycerol gradient and native gel. The 44.4 kDa polypeptide was chiefly incorporated into an ~1 MDa (30S) complex that extended into heavier fractions (35S-50S); a minor ~190 kDa MERS1-containing particle was also detected (Fig. 3A, left panel). Notably, lysate pre-treatment with RNase I released MERS1 from high molecular mass complex as a discrete ~160 kDa particle (Fig. 3A, middle panel). We also noticed that the high molecular mass MERS1 complex closely resembles patterns displayed by GRBC1/2 proteins (Fig. 3A, right panel). These proteins are responsible for gRNA stabilization (20) and belong to the gRNA binding module (GRBC) within the RNA editing substrate binding complex (RESC). Two other RESC modules, RNA editing mediator (REMC) and polyadenylation mediator (PAMC), engage the U-insertion/deletion mRNA editing core (RECC) and polyadenylation (KPAP) complexes, respectively (21).

**Fig. 3.**
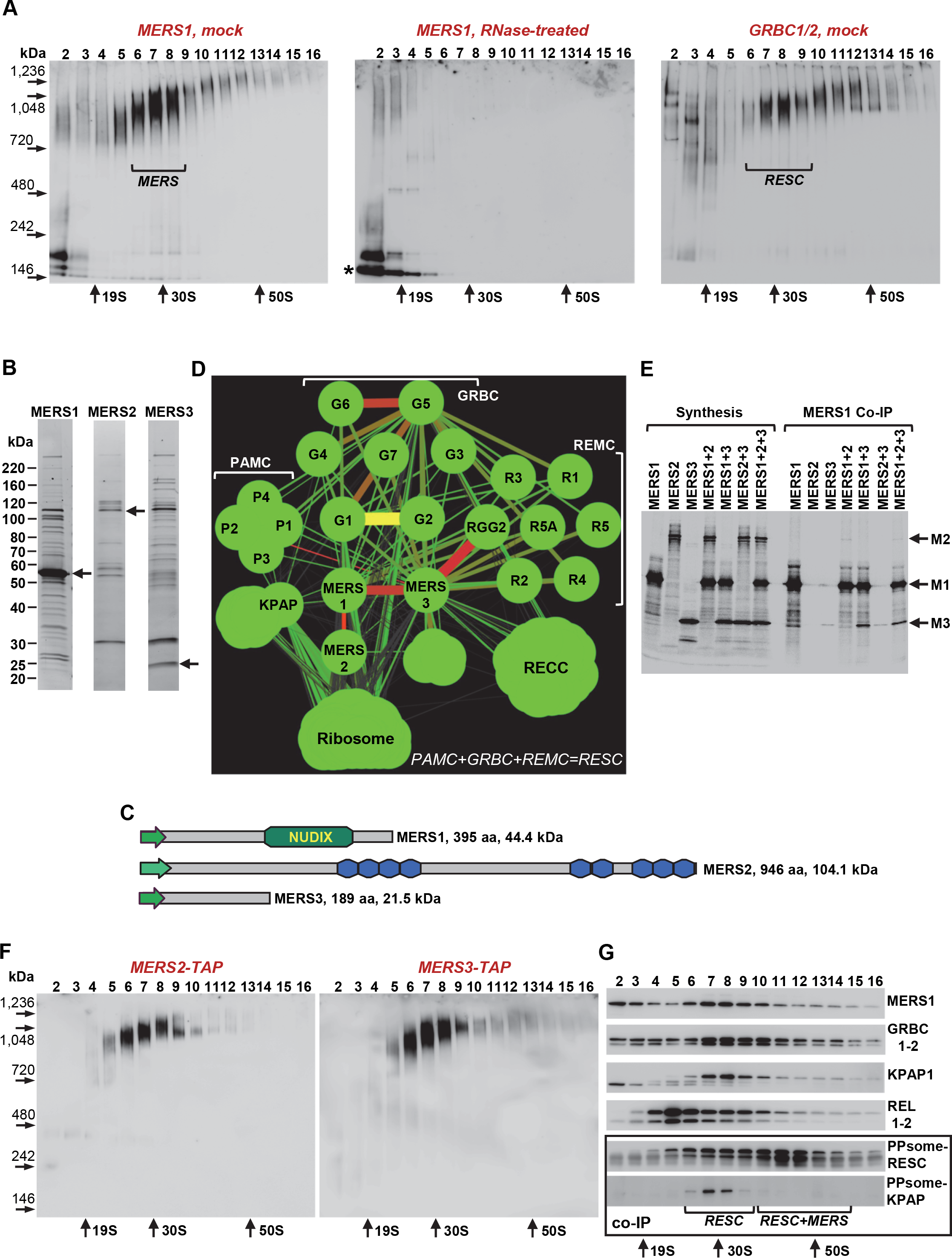
PPsome interacts with mRNA processing complexes. (A) MERS1 interacts with a high molecular mass complex via RNA. Mock-and RNase-treated cell lysates were fractionated on 10 – 30% glycerol gradients, and each fraction was further resolved on native 3 – 12% Bis-Tris acrylamide gel. MERS1 complex was detected by immunoblotting with polyclonal antibodies against recombinant protein (left and middle panels). MERS1 particle released by RNase treatment is marked by asterisk. The RNA editing substrate binding complex (RESC) was visualized with anti-GRBC1/2 antibodies in mock-treated extract only, right panel. Positions of molecular mass and sedimentation markers are indicated. (B) Tandem affinity purification of PPsome components. Purified MERS1, 2 and 3 fractions were separated on SDS page and stained with Sypro Ruby. Bait proteins are indicated by arrows. (C) Domain organization of major PPsome components. MERS1 hydrolase domain and pentatricopeptide repeats in MERS2 are diagrammed. MERS3 lacks any discernable motifs. Green arrows show mitochondrial targeting peptides. (D) Interactions network of MERS1, 2 and 3, and polyadenylation (KPAP), RNA editing substrate binding complex (RESC), RNA editing core complex (RECC), and the ribosome. The RESC complex is composed of guide RNA binding (<u>G</u>RBC), RNA editing mediator (<u>R</u>EMC) and polyadenylation mediator (<u>P</u>AMC) modules (21). Their subunits are marked as G, R and P, respectively. The network was generated in Cytoscape from bait-prey pairs in which the prey protein was identified by five or more unique peptides. The edge thickness and color correlate with normalized spectral abundance factor (NSAF) value. (E) *In vitro* reconstitution of the PPsome. Synthesis: Individual proteins, or their combinations, were synthesized in a coupled transcription-translation reticulocyte system supplemented with [^35^S] methionine. MERS1 Co-IP: Immunoprecipitations were performed with immobilized anti-MERS1 polyclonal antibody. Co-precipitated proteins were eluted from antibody-coated beads, separated on 8 – 16% SDS PAGE, and exposed to phosphor storage screen. Positions of PPsome subunits are indicated by arrows as M1, M2 and M3. (F) Complex association of MERS2 and MERS3 subunits. Extracts from cells expressing respective C-terminally TAP-tagged proteins were fractionated as in (A). Fusion proteins were detected by immunoblotting with antibody against calmodulin binding peptide tag. (G) Interactions between PPsome, RESC and KPAP (polyadenylation) complexes. Gradient fractions were separated on denaturing PAGE and probed for MERS1, GRBC1/2 and KPAP1 poly(A) polymerase. RNA editing core complex (RECC) was detected by self-adenylation of REL1 and REL2 ligases in the presence of [α-^32^P]ATP. RESC and KPAP1 were visualized in samples immunoprecipitated with anti-MERS1 antibody (framed panels). Note that PPsome-RESC co-complex interactions extend into the 40S-50S segment.

To gain a higher-resolution view, we performed LC-MS/MS analysis of tandem affinity-purified MERS1, and two proteins that were most abundant in the MERS1 fraction. These were the pentatricopeptide repeat-containing (PPR) protein termed MERS2 (Tb11.02.5120) and MERS3 polypeptide lacking any discernible motifs (Tb927.10.7910) (Fig. 3B, C). The established components of the RESC (GRBC1 and GRBC5), RNA editing core (RET2 TUTase) and polyadenylation (KPAP1) complexes were also purified along with small (S17) and large (L3) ribosomal subunits (Dataset S2). An interaction network built on normalized spectral abundance factors (25), predicted that MERS1 interacts with a MERS2, and connects to RESC through a MERS3 (Fig. 3D). To validate the MERS1-MERS2 and MERS1-MERS3 interactions, we performed co-immunoprecipitations from reticulocyte lysates programmed for synthesis of binary or ternary combinations (Fig. 3E). When accounted for methionine residues, MERS1 and MERS3 formed a stoichiometric complex; MERS1-MERS2 binding was apparently less stable, but still detectable after stringent washes. Furthermore, Fig. 3F shows that TAP-tagged MERS2 and MERS3 are confined to a high molecular mass complex matching the size of MERS1-contaning particles. Based on these results, we designated MERS1, 2 and 3 complex as the mitochondrial 5′ <u>p</u>yro<u>p</u>hosphohydrolase (PPsome). The co-complex interactions between 5′ PPsome, RESC and KPAP complexes were confirmed by MERS1, GRBC1/2 and KPAP1 poly(A) polymerase co-immunoprecipitation along glycerol gradient (Fig. 3G). Collectively, fractionation and reconstitution studies indicate that the PPsome engages in RNA-mediated interactions with the RNA editing substrate binding and polyadenylation complexes.

### PPsome Binding to 5′ Termini Stabilizes mRNAs and 9S rRNA

In *T. brucei*, unedited and edited mRNAs exist in two forms distinguished by 3′ modification patterns: short A-tail (20-25 nt) and bi-modal 200-300 nt-long A/U-tail in which short A-tail is extended into A/U heteropolymer (26–28). Pre-edited mRNAs possess only short A-tails while rRNAs and gRNAs are uridylated. Polyadenylation plays a key role in mRNA stabilization (19, 27, 29), while contribution of the 5′ processing remains unexplored. MERS1 RNAi knockdown triggered rapid decline of pre-edited and edited RPS12 (Fig. 4A), and unedited CO1 mRNA (Fig. 4B). Small ribosomal 9S RNA was also moderately downregulated (Fig. 4C), while 12S rRNA and guide RNAs (Fig. 4D) remained unchanged. To test whether accelerated mRNA and 9S rRNA decay in MERS1 RNAi cells may account for the observed changes in steady-state levels, we performed real-time decay assay. Here, MERS1 is depleted by RNAi, and transcription is then blocked with ethidium bromide and Actinomycin D (17). Decay kinetics demonstrated that MERS1 knockdown causes moderate stabilization of short-and long-tailed edited mRNA forms, but accelerated degradation of pre-edited mRNA (Fig. S2). Hence, the decline of pre-edited precursor causes the loss of edited mRNA. The ribosomal RNA decay kinetics were consistent with drop in 9S, but not 12S RNA abundance. Thus, MERS1 is essential for stability of most RNAs transcribed from the maxicircle but is dispensable for 12S rRNA and guide RNA.

**Fig. 4.**
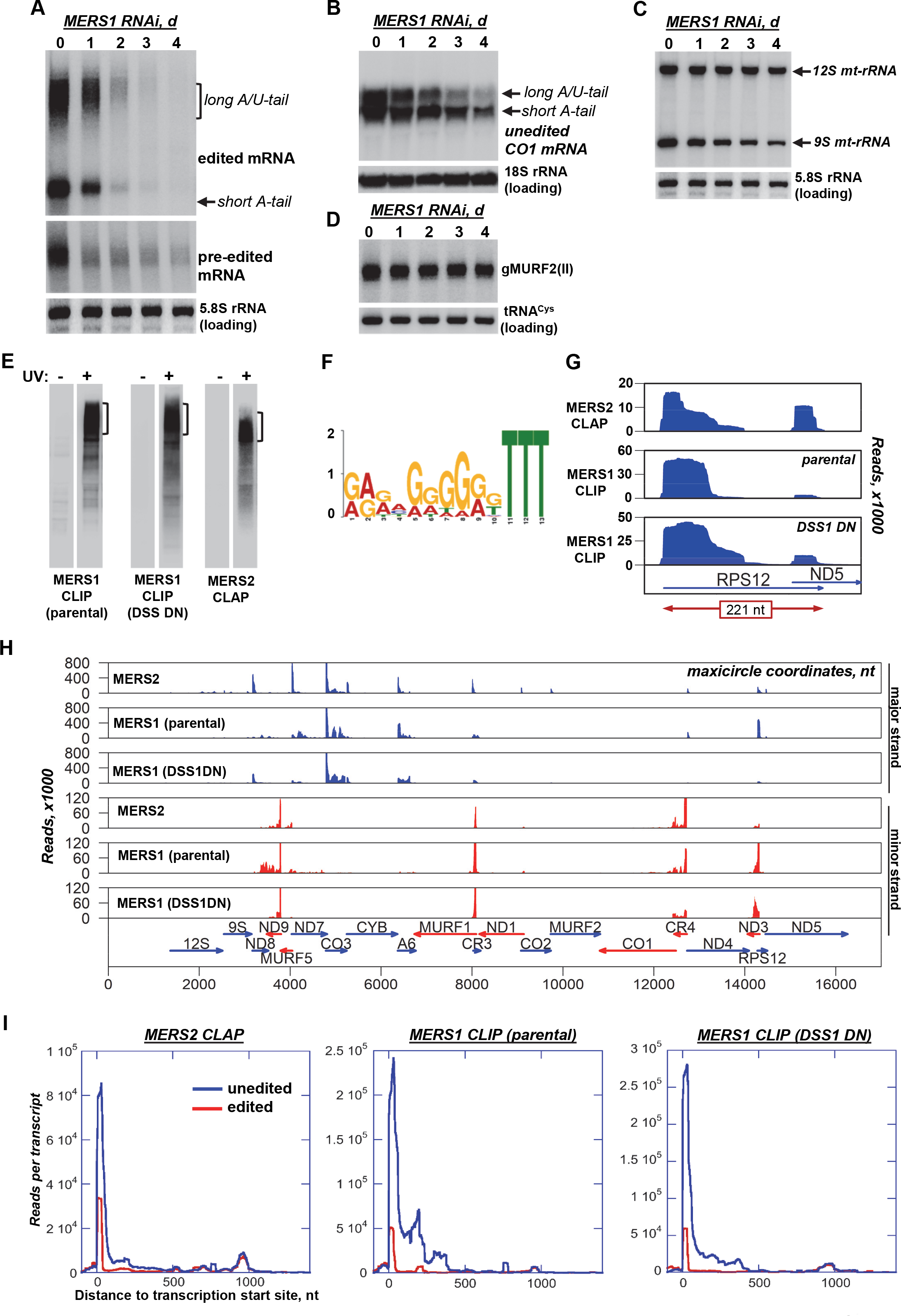
MERS1 is essential for mRNA and 9S rRNA stability. (A) MERS1 RNAi knockdown impact on pan-edited mRNA. Edited and pre-edited forms of representative RPS12 mRNA were analyzed by Northern blotting. The short A-tailed and long A/U-tailed edited mRNA populations are indicated. RNAi was induced by adding tetracycline. MERS1 downregulation was verified by immunoblotting (Fig. 5D). (B) MERS1 knockdown effects on unedited mRNAs was assessed by Northern blotting. (C) Ribosomal RNA Northern blotting in MERS1 RNAi cells. (D) Guide RNA Northern blotting in MERS1 RNAi cells. (E) Isolation of UV-induced MERS1-and MERS2-RNA crosslinks. CLIP: MERS1 was immunoaffinity-purified with polyclonal antibody from parental and DSS1 DN cells. CLAP: TAP-tagged MERS2 was purified by affinity pulldown with IgG-coated magnetic beads. Parasites were mock-treated (−) and UV-irradiated (+). RNA was fragmented by RNase A and T1, radiolabeled, released from the crosslink (areas indicated by brackets), and sequenced. (F) MERS2 *in vivo* binding motif. The MEME algorithm was applied to predict the sequences enriched in MERS2-crosslinked RNA. (G) MERS2 CLAP and MERS1 CLIP (from parental and DSS1 DN cells) reads were aligned to a representative maxicircle region with overlapping mRNAs encoded on the same strand. (H) PPsome binding sites in the maxicircle. MERS1 CLIP and MERS2 CLAP reads from parental and DSS1DN cell lines were mapped to the gene-containing region. Annotated mitochondrial transcripts, read count scales and maxicircle coordinates are indicated. (I) Composite distribution of PPsome complex binding sites in mitochondrial mRNAs. The reads were aligned to unedited (blue) and fully-edited sequences (red). Read counts located 1000 nt downstream and 100 nt upstream of the 5′-end in each transcript were collected in 1 nt bin. The average coverage across all maxicircle genes was plotted.

**Fig. 5.**
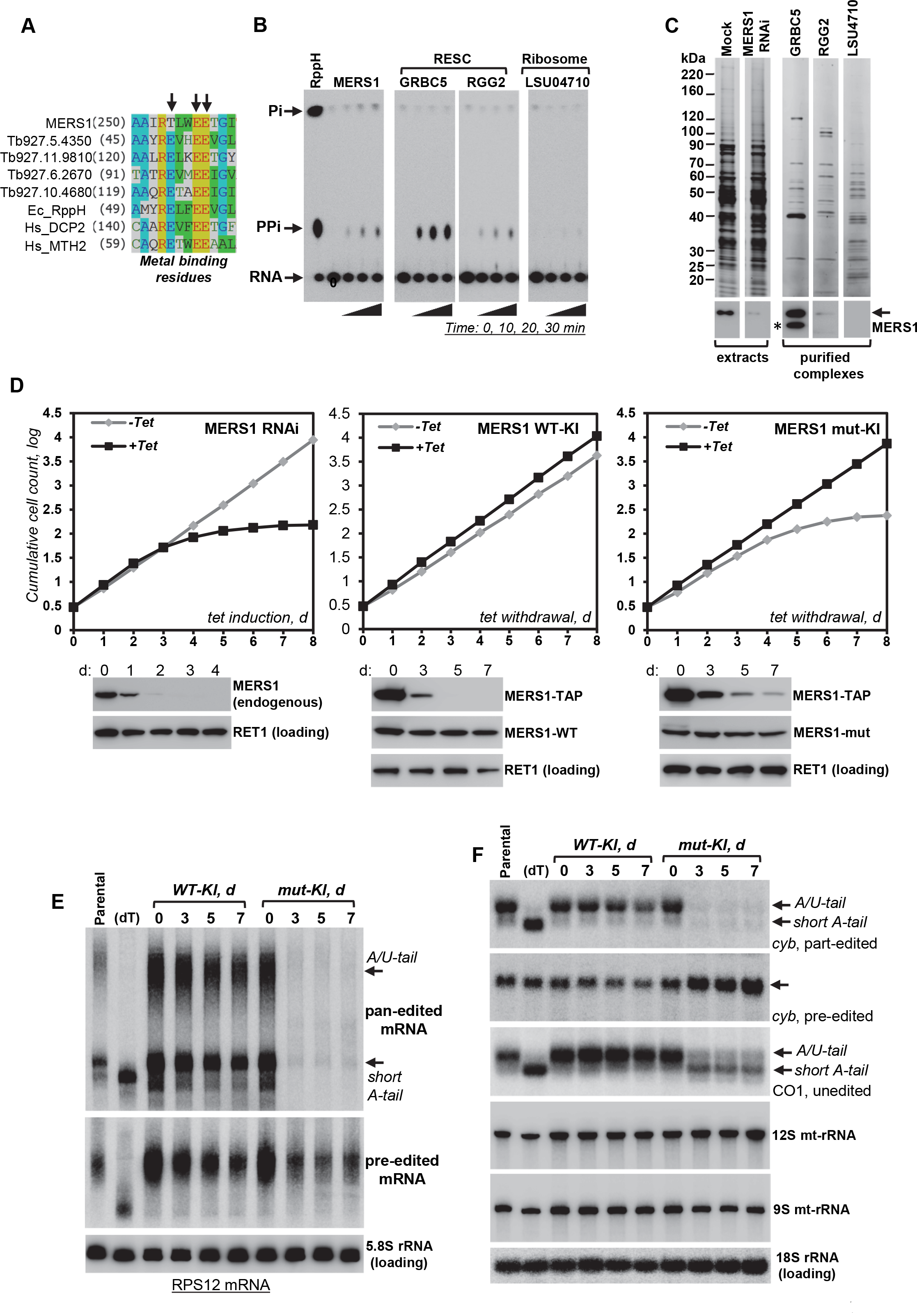
RESC complex stimulates MERS1 activity. (A) Partial alignment of NUDIX motifs from trypanosomal, bacterial and human pyrophosphohydrolases. EC_RPPH; *Escherichia coli* RppH (WP_088540307.1), HS_DCP2; *Homo sapiens* Dcp2 (NP_001229306.1), HS_MTH2; *H. sapiens* Mth2 (NP_060753.1). Metal binding acidic residues are shown by arrows. (B) Pyrophosphohydrolase activity of purified complexes. The MERS1 and RESC complexes, and the ribosome were affinity-purified via indicated subunits and incubated with 5′ [γ-^32^P]-labeled RNA. Reaction products were separated by thin-layer chromatography along with those produced by RppH NUDIX hydrolase from *E. coli*. (C) MERS1 relative abundance in purified complexes. Mock-treated and RNAi knockdown cells were analyzed along with tandem affinity purified RESC and the ribosome. Cross-reactivity of polyclonal antibodies raised against 6-His tagged recombinant MERS1 with likewise tagged GRBC5 bait is shown by an asterisk. (D) Cell growth kinetics of MERS1 RNAi and conditional knock-in (KI) cell lines. RNAi was induced with tetracycline to downregulate endogenous MERS1. In the knock-in, the drug was withdrawn to suppress conditional MERS1-TAP expression in KI cells. One endogenous allele constitutively expressed functional (WT-KI), or inactive (mut-KI), MERS1 proteins while the other allele was disrupted (Fig. S4A). RNAi repression, and conditional (MERS1-TAP) and constitutive (MERS1-WT and MERS1-mut) expression were verified by Western blotting. (E) Effects of MERS1 enzymatic activity loss on pan-edited mRNA. Pre-edited and edited forms of RPS12 mRNA were detected by Northern blotting. The short A-tailed and long A/U-tailed edited mRNA populations are indicated. (dT), RNA was treated with RNase H in the presence of oligo(dT) 20-mer to remove short A-tails and long A/U-tails. (F) Effects of MERS1 enzymatic activity loss on moderately-edited *cyb*, and unedited CO1 mRNAs, and 9S and 12S rRNAs.

Messenger RNA stabilization by the polyadenylation complex has been linked to Kinetoplast Polyadenylation Factor 3 (KPAF3) binding near the 3′-end and A-tailing by KPAP1. These events inhibit mRNA degradation by DSS1 exonuclease (19, 27). To distinguish whether 5′ PPsome binds near 5′ triphosphate or, similarly to polyadenylation complex, to the 3′-end, we identified MERS1 and MERS2 binding sites by *in vivo* UV crosslinking (Fig. 4E). The highly similar sequences derived from MERS1-CLIP and MERS2-CLAP (Pearson correlation score 0.774, P-value = 2.2×10^−16^) indicate their binding to the same purine-rich sites, with a strong bias for three uridines at the 3′-end (Fig. 4F). Importantly, PPsome binds chiefly to the 5′ extremity of annotated mRNAs (Fig. 4G, H), while less than 1% of MERS1 or MERS2 crosslinks could be mapped exclusively to minicircle-encoded gRNAs (Dataset S3). Finally, CLIP experiments performed in DSS1 DN cells showed that PPsome binding patterns remain unaltered (Fig. 4H, I) notwithstanding accumulation of 3′ extended precursors upon DSS1 repression (19). Together, these results implicate the PPsome in recognizing specific sequences adjacent to transcription-generated 5′-end and preventing mRNA degradation by DSS1 exonuclease.

### RNA Editing Substrate Binding Complex Stimulates PPsome Activity

NUDIX-like activity would be expected to remove pyrophosphate, or sequentially hydrolyze γ-and β-phosphates from the 5′-end of a primary transcript (30). However, the recombinant MERS1 purified from bacteria was inactive with a substrate derived from PPsome binding site in ND7 pre-mRNA. Partial multiple sequence alignment of the “NUDIX box” from trypanosomal, bacterial and human enzymes identified replacement of a conserved catalytic glutamic acid residue by a threonine in MERS1 (Fig. 5A). This substitution is conserved in MERS1 proteins from other kinetoplastids (not shown) and may account for the compromised activity. To establish whether complex association is required to activate MERS1, and to determine the nature of the leaving group, we performed an enzymatic assay with affinity-purified PPsome. The time-dependent accumulation of pyrophosphate demonstrated that MERS1 indeed possesses the expected pyrophosphohydrolase activity (Fig. 5B). To investigate functional significance of PPsome interaction with the RNA editing substrate binding complex (Fig. 3), we tested the hydrolase activity in the RESC complex variants purified via GRBC5 and RGG2 subunits, and large ribosomal subunit as control. Remarkably, the RESC-bound PPsome displayed higher rate of activity (Fig. 5B), which correlated with MERS1 abundance in GRBC5 and RGG2 preparations (Fig. 5C). Adjusted for MERS1’s relative abundance (Dataset S2), association with RESC stimulated PPsome activity by ~100-fold.

MERS1 downregulation by RNAi led to mRNA and 9S rRNA decline (Fig.s 4 and S3) and cell death (Fig. 5D) indicating that PPsome binding to the 5′-end is essential for mRNA and 9S rRNA stabilization (Fig. S2). Knockdowns of *MERS2* and *MERS3* expression showed that the encoded proteins are also essential for normal growth and that their loss affects several edited mRNAs (Fig. S3). However, protein depletion may cause a phenotype due to loss of intrinsic activity, or by preventing assembly of a functional complex. To verify MERS1 enzymatic identity and to address the functional significance of tri-to monophosphate conversion, we generated procyclic cell lines for conditional MERS1 knock-in. In these backgrounds, one allele is disrupted, then a *tet*-repressor-controlled TAP-tagged copy of a functional gene is introduced into the rRNA locus and kept actively expressed by maintaining drug in the media. The second allele is then replaced with a cassette expressing either a functional or inactive (E257A, E258A) gene (Fig. S4A). Upon tetracycline withdrawal, the *tet* repressor blocks the ectopic expression. As MERS1-TAP gradually declines, the parasite’s survival relies on the functionality of the mutated MERS1 expressed from an endogenous allele. In these settings, monoallelic MERS1 expression was sufficient to sustain cell division, while the active site mutations led to a pronounced growth inhibition phenotype (Fig. 5D).

Northern blotting of pan-edited (RPS12), moderately-edited (*cyb*), and unedited (CO1) mRNAs along with rRNAs (Fig. 5E, F), and qRT-PCR (Fig. S4B) confirmed the virtually identical impacts of mutated MERS1 monoallelic expression and RNAi knockdown of an endogenous protein (Fig. 4). In addition, instructive differences were observed between pan-and moderately-edited RNAs. In pan-edited RPS12 mRNA, editing events occur within ~20 nt from the polyadenylation site and expand toward the 5′-end nearly doubling mRNA length in the process (31, 32). In moderately-edited *cyb* mRNA, 34 uridines are inserted adjacent to the MERS1 binding site at the 5′-end. In contrast to RPS12 mRNA, pre-edited *cyb* transcript accumulated, while the edited form declined. Thus, PPsome binding may reciprocally stimulate editing in the proximity to the 5′-end by recruiting the RESC complex.

### 5′ Pyrophosphate Hydrolysis and 3′ Adenylation Are Independent Events

The mechanism of 5′-end formation described above resolves the long-standing observation of monophosphorylated mRNAs and rRNAs and vacates the hypothetical endonuclease involvement. The scarcity of PPsome recognition sites in guide RNAs also explains 5′3P status of these molecules. The pyrophosphate removal and PPsome deposition at the 5′-end apparently block mature mRNA degradation by DSS1 3′-5′ exonuclease, but not 3′-5′ precursor processing by the same activity. It follows that polyadenylation factors binding at the properly trimmed 3′ end and polyadenylation by KPAP1 are required (19, 27), but not sufficient for mRNA stabilization. Based on a RESC-mediated link between the PPsome and polyadenylation complex (Fig. 3), we hypothesized that MERS1-dependent mRNA stabilization is contingent upon accurate 3′-end formation and polyadenylation. Mitochondrial mRNAs display temporally separated and functionally distinct 3′ modifications: Short A-tails (~20 nt) are added prior to editing, and then extended into a long A/U-heteropolymers (~200 nt) upon completion of editing. Irrespective of editing history, the A-tail stabilizes edited and unedited mRNAs, while the A/U-tail manifests translation-competent mRNAs capable of binding to the ribosome (28). To investigate whether MERS1 repression compromises mRNA stability by interfering with 3′ adenylation, 3′ RACE RNA-Seq libraries were constructed to map unmodified, adenylated and uridylated termini in parental, MERS1 RNAi, and DSS1 DN cell lines. As expected, DSS1 repression led to a decline in adenylated 3′-ends and an increase in unmodified precursors. Conversely, MERS1 RNAi exerted only minor global effects on mRNA polyadenylation (Fig. 6A). Positioning (Fig. 6B) and nucleotide composition (Fig. 6C) of functional A-tails and cryptic (unmodified, mixed tails and U-tails) 3′ modifications also remained unaltered in pan-edited RPS12 mRNA. A summary of mature 5′ and 3′ termini determined by RACE experiments is presented in Dataset S4. We conclude that MERS1 knockdown does not affect 3′ adenylation. Detection of uniform 5′P termini in KPAP1 poly(A) polymerase (27) and DSS1 (19) knockdowns further establishes that blocked 3′ adenylation or trimming do not impact PPsome activity. Although the 5′ and 3′ processing events occur independently, the RESC-mediated interaction between PPsome and polyadenylation complex occupying the respective mature termini apparently protects RNA against degradation by DSS1 exonuclease.

**Fig. 6.**
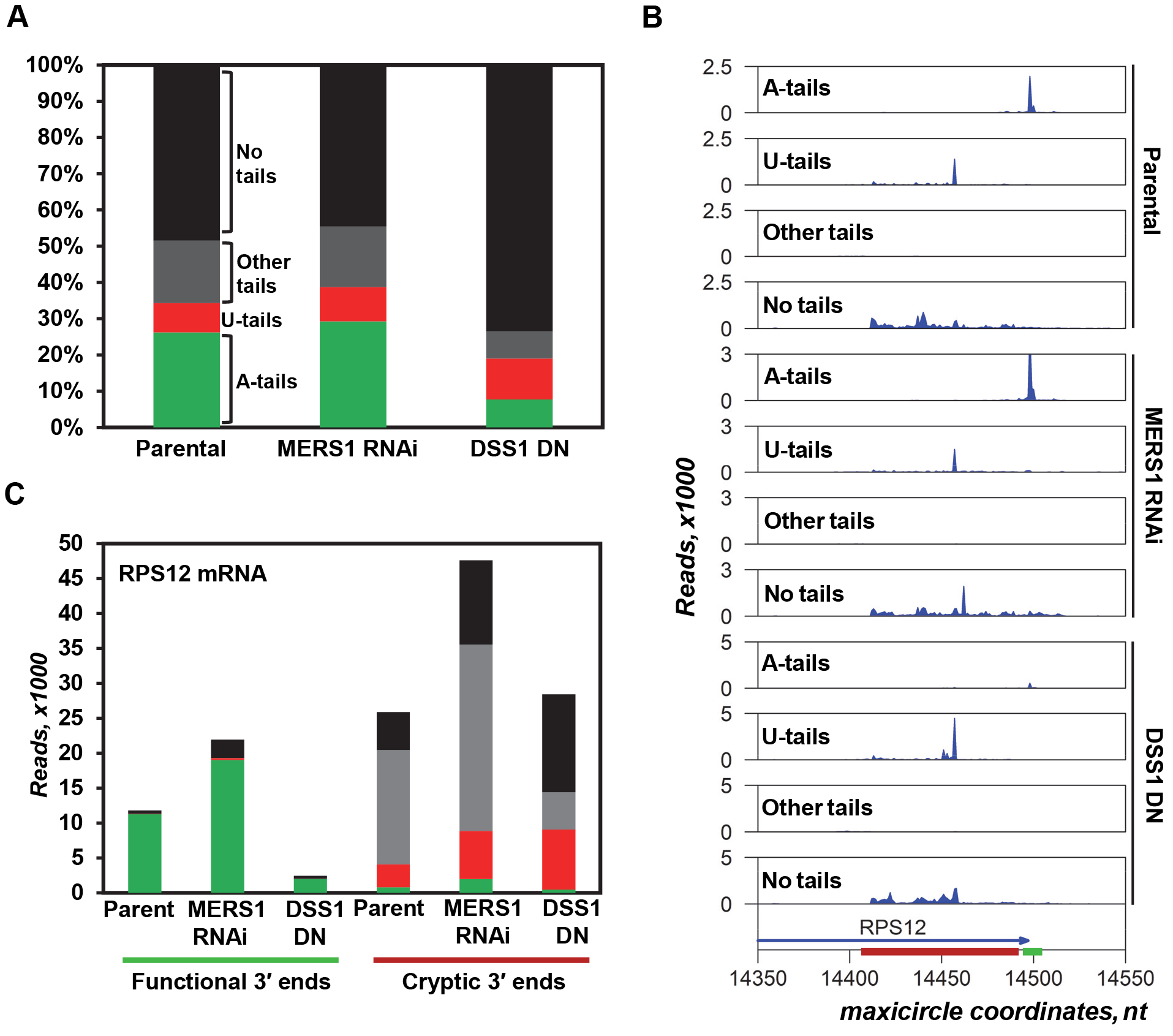
Adenylation is unaffected by MERS1 repression. (A) Relative global abundance of mRNA 3′ modifications. RNA linker ligation-based 3′ RACE was performed on parental, MERS1 RNAi and DSS1 DN cells. The modifications were classified as A-tail (>90% As), U-tail (>90% Us), other (no nucleotide constitutes more than 90%), and unmodified. Read counts were normalized to synthetic RNA spike. (B) RPS12 pre-mRNA processing variants clustered by tail type. The last encoded nucleotide of mRNA bearing the same tail was mapped to the *RPS12* gene (blue arrow). Read scale and maxicircle coordinates are indicated. Functional mRNA 3′ UTR containing stop codon is underlined by a green bar; red bar shows truncated variants. Read counts were normalized to synthetic spike RNA. (C) Tail composition of functional and truncated RPS12 mRNA 3′ ends. Color code as in (A).

### Antisense Transcription Defines Mature 3′-end of Maxicircle-encoded RNAs

The accurate precursor trimming by DSS1 is required for the Kinetoplast Polyadenylation Factor 3 (KPAF3) binding to 3′-end and KPAP1 poly(A) polymerase recruitment (19). The ensuing A-tailing and bridging between polyadenylation complex and the PPsome likely lead to mRNA stabilization. However, KPAF3 binding is incapable of stopping 3′-5′ degradation at a precise position (19), which emphasizes the central role of 3′-end definition in mRNA biogenesis and stabilization. To determine the mechanism of mRNA 3′-end formation, we hypothesized that antisense transcription near the annotated mRNA 3′-end yields non-coding RNAs capable of impeding DSS1 activity. In this scenario, the 5′-end of an antisense RNA would define positioning of the mRNA 3′-end. Rapid degradation of antisense transcripts would be expected to liberate pre-mRNA for further processing by internal editing. The 5′ RACE RNA-Seq library was constructed from mock-and polyphosphatase-treated RNA to identify potential antisense transcription initiation sites near annotated mRNA 3′ termini in parental, MERS1 RNAi and DSS1 DN backgrounds. By juxtaposing 3′ RACE-derived mRNA polyadenylation and 5′ RACE-generated antisense transcription initiation composite profiles, we detected consistent initiation signals extending inward from canonical polyadenylation sites (Fig. 7A). Remarkably, strong antisense initiation signals were also observed near previously mapped truncated mRNAs 3′ termini, which are often uridylated (−70 to −80 region, (19)). To confirm non-coding RNA synthesis by antisense transcription, we performed Northern blotting to visualize molecules that are complementary to RPS12 (major strand) and MURF5 (minor strand) pre-mRNAs (Fig. 7B). The detected transcripts did not correspond to any annotated mRNAs transcribed from the same strand and location, and apparently lacked 3′ A-tails. Furthermore, in contrast to canonical mRNAs (Fig. 4), the non-coding RNAs were upregulated in both MERS1 RNAi and DSS1 DN cells. While antisense accumulation is expected in DSS1 DN background with inhibited processing exonuclease, a similar build up in MERS1 knockdown implicates the PPsome in destabilizing the non-adenylated antisense transcripts. In agreement with the 3′RACE results, adenylation is apparently required for RNA stabilization by the PPsome.

**Fig. 7.**
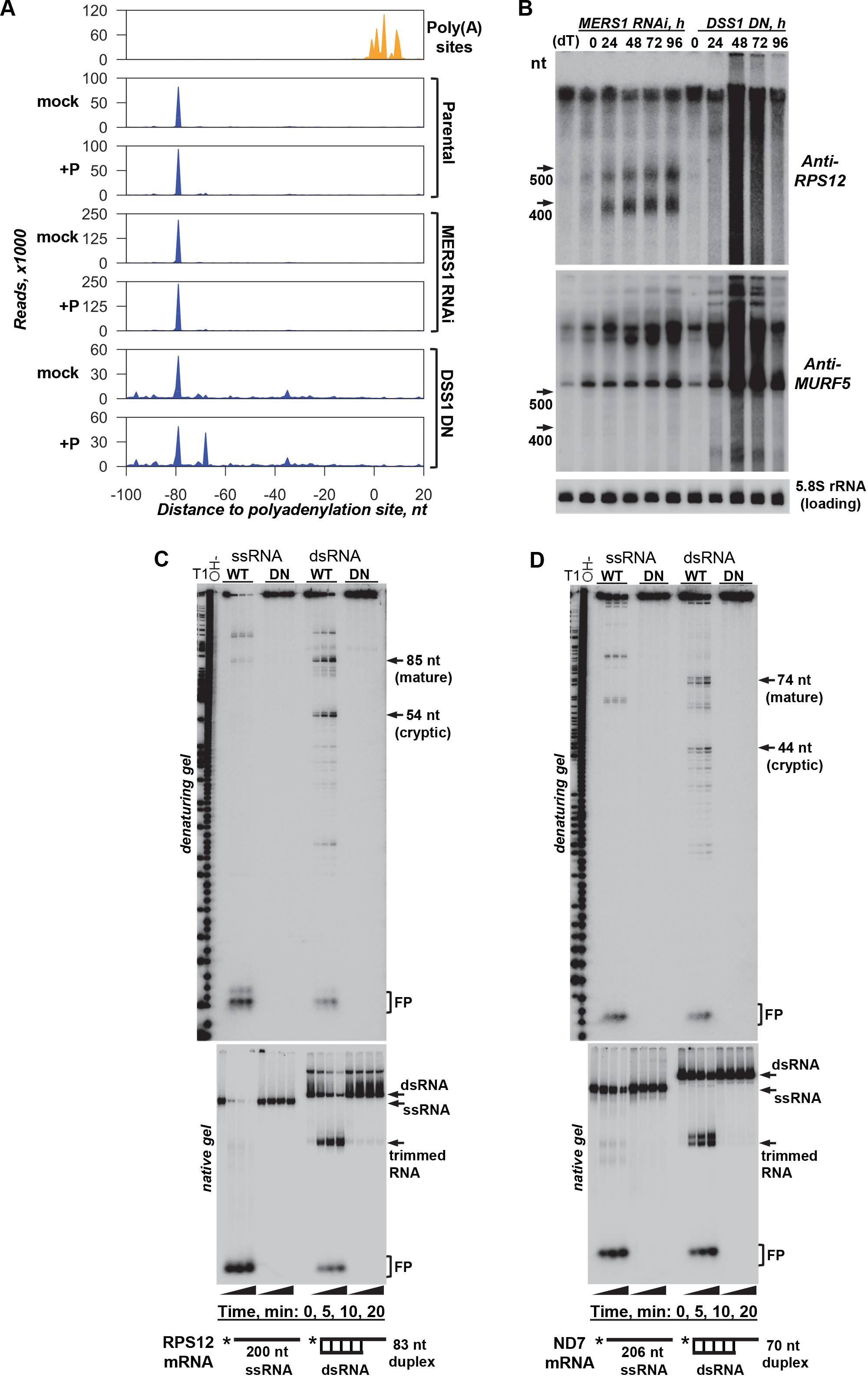
Antisense transcription defines mRNA 3′ end by blocking DSS1 exonuclease. (A) Mapping 5′ termini of antisense transcripts. A 5′ RACE was performed in the parental, MERS1 RNAi and DSS1 DN cells. RNA was treated with 5′ polyphosphatase to capture mono-and triphosphorylated transcripts (+P), or mock treated. Positions of canonical mRNA adenylation sites are shown in the top panel. The 5′ RACE reads for antisense RNAs were aligned to maxicircle sequences. Read counts located 100 nt downstream and upstream of the mapped polyadenylation site in each transcript were collected in a 1 nt bin. Composite distribution of antisense RNA 5′ ends within annotated mRNA boundaries is shown by summation of coverage across all genes. The 3′ RACE-defined polyadenylation site is set as zero. Read scale and maxicircle coordinates are indicated. (B) Detection of non-coding RNAs transcribed as antisense to major strand-encoded RPS12 and MURF5 pre-mRNAs. Total RNA from MERS1 RNAi and DSS1 DN cells was analyzed by Northern blotting. (dT), RNA was treated with RNase H in the presence of oligo(dT) 20-mer to eliminate A-tails. Note unaltered migration patterns in RNase H/oligo(dT) sample. (C) Antisense RNA-controlled 3′ end definition *in vitro*. Active (WT) and inactive (DN) DSS1 exonuclease variants were isolated from mitochondrial fraction by tandem affinity purification. Reactions with 5′ radiolabeled single-stranded (ss) RPS12 mRNA fragment, or pre-assembled partially double-stranded (ds) RNAs, were terminated by adding Proteinase K. Products were resolved on polyacrylamide denaturing (upper panel) or native (lower panel) gels. FP, final degradation products (4-5 nt). (D) Same reactions as in (C) were performed with ND7 mRNA fragment.

To test whether antisense non-coding RNA can impede highly-processive DSS1 exonuclease activity, we have reconstituted the mRNA 3′ processing reaction *in vitro*. Synthetic 5′ radiolabeled RNAs that resemble 3′ regions of RPS12 and ND7 pre-mRNAs were hybridized with data-supported antisense RNA fragments and incubated with affinity-purified active (WT) and inactive (DN) DSS1 3′-5′ exonuclease (Fig. 7C, D, upper panels). Both single-stranded fragments were processively degraded to 4-5 nucleotides (FP) with minor amounts of abortive products, which is consistent with an RNase II-like DSS1 activity (16). Introducing partially double-stranded RNA into the reaction induced a strong and precise pausing 3-4 nt upstream of the antisense RNA 5′-end, but also ~30 nt shorter RNAs that resemble cryptic internal stops observed *in vivo* (Fig. 6B). To verify formation of a duplex with a trimmed 3′ overhang, the reaction products were separated on native gel under conditions that retain all degradation products (Fig. 7C, D, lower panels). The DSS1 pausing before sDataset duplex regions appears to be stochastic: the RNA hydrolysis-driven unwinding activity proceeds with some frequency into the double-stranded region or degrades the entire RNA to short oligonucleotides. The former pattern would be consistent with detection of truncated mRNA 3′-ends, while the latter explains accumulation of mRNA precursors at significantly higher levels than mature mRNAs (17, 19).

## Discussion

The mitochondrion undergoes dramatic changes in function, size and gene expression during digenetic life cycle of *Trypanosoma brucei*. The developmental variations in abundance, the 3′ modification state and the extent of editing have been documented for most mitochondrial mRNAs (33), but few of these factors correlate with expected requirements for a specific protein at a particular life cycle stage. Incongruously, little is known about signaling mechanisms that align mitochondrial function with rapidly changing environment during transmission and infection. Major advances in understanding mRNA editing, polyadenylation and translation processes (31, 34) have left the decades-old notion of unregulated multicistronic transcription unperturbed. Likewise, the primary RNA cleavage by a cryptic endonuclease was assumed to produce monocistronic substrates for 3′ adenylation and editing. Although this scenario explains monophosphorylated 5′ and homogenous 3′ termini, the endonuclease identity and specificity determinants remained unsolved. Inherently, the endonuclease model negates transcriptional control of individual gene activity and presumes that adenylation and editing ultimately dictate the steady-state levels of translation-competent mRNAs. By combining *in vivo* nucleic acid – protein crosslinking, genetic, proteomic, and *in vitro* reconstitution studies of mitochondrial RNA polymerase, and mRNA 5′ and 3′ processing, and editing complexes, we show that maxicircle genes are individually transcribed as 3′ extended precursors. We demonstrate that mRNA and rRNA 5′ termini are set by transcription initiation and dephosphorylated by the MERS1 NUDIX hydrolase. Acting as subunit of the 5′ pyrophosphohydrolase (PPsome), MERS1 is stimulated by RNA-mediated association with the RNA editing substrate binding complex (RESC). It seems likely that RESC recruitment serves as a two-pronged quality check point to: 1) Ensure that only target-bound MERS1 is catalytically active, and 2) Verify that RESC-bound pre-mRNA is properly adenylated, and, therefore, competent for internal editing. These conclusions are also supported by stronger downregulation of edited mRNAs versus their pre-edited counterparts, such as *cyb* mRNA (Fig. 5F), or unedited transcripts. The specificity, however, comes at a cost. The low efficiency of MERS1-catalyzed reaction may be responsible for rapid decay of mRNA precursors in contrast to structured rRNAs, which are less affected by MERS1 repression. Indeed, MTRNAP maxicircle occupancy indicates that mRNA genes are populated by RNA polymerase at 2 – 3-fold higher level while mature rRNAs are more abundant than mRNAs by 10 – 100-fold.

PPsome *in vivo* RNA binding sites are predominantly located near the mRNA 5′ termini, but it seems likely that MERS2 pentatricopeptide repeat subunit is responsible for RNA recognition and enabling MERS1 activity. Although maxicircles constitute only about 5% of kDNA mass (35), their transcripts accounts for more than 65% of PPsome binding sites, while only ~1% belong to the highly diverse and abundant minicircle-encoded gRNAs. This correlation indicates that gRNAs are not recognized by the PPsome, and explains why this class of mitochondrial RNA maintains transcription-incorporated 5′ triphosphate. Conversely, PPsome-bound primary 5′-ends of maxicircle transcripts are converted into monophosphate form. The essentiality of MERS1-catalyzed pyrophosphate hydrolysis for mRNA and rRNA maintenance underscores the fundamentally different mechanisms that stabilize maxicircle-encoded mRNAs and minicircle-encoded gRNAs. The mature 3′ uridylated gRNAs are directly bound to the guide RNA binding complex (GRBC (17, 20)), a discrete module within the larger RESC complex (Fig. 3, (21, 36)). Initially defined as the RNA binding component of the RNA editing holoenzyme, RESC also interacts with the polyadenylation complex, consisting of KPAP1 poly(A) polymerase (27), and at least two PPR polyadenylation factors with distinct functions. KPAF3 PPR binds 3′-end to stabilize mRNA and to stimulate short A-tail addition prior to editing (19), while KPAF1/2 dimer induces post-editing A/U-tailing to activate translation (28). The KPAF3 binding and ensuing A-tailing are necessary, but not sufficient determinants of mRNA stability. Our study demonstrates a critical role of 5′ PPsome in protecting mRNA against degradation by DSS1 exonuclease, which is responsible for both 3′ processing and decay of all mitochondrial RNA species (16, 19). The RESC-mediated contacts between PPsome and polyadenylation complexes strongly suggest that a proximity of monophosphorylated 5′ and adenylated 3′-ends may be an essential mRNA stabilization element. The interaction between PPsome and the ribosome also hints at 5′ “protein cap” involvement in translation, but this inference requires further investigation.

Considering gene-specific transcription initiation, the RNAs spanning gene boundaries (12, 37) apparently represent 3′ heterogeneous precursors that intrude into downstream coding sequences, but are inevitably trimmed back by DSS1 (19). Therefore, the 3′-end processing pathway should include a mechanism by which the highly processive 3′-5′ precursor degradation is blocked at a specific point prior to mRNA adenylation or rRNA uridylation. To that end, we have identified ubiquitous antisense transcripts that initiate near the functional mRNA 3′-end and reconstituted the 3′-5′ degradation pausing *in vitro*. In this stochastic event, the 5′-end of the antisense non-coding RNA dictates the position of mRNA’s 3′-end and generates a substrate for the polyadenylation complex. In summary, we demonstrated that mitochondrial pre-mRNAs and rRNAs are transcribed individually and revealed the mechanisms by which 5′ and 3′ termini are produced, and mature mRNAs are stabilized. It is now conceivable that transcriptional control, in addition to internal editing and 3′ adenylation mechanisms (17, 19, 27), plays a significant role in developmental regulation of mitochondrial gene expression.

## Methods

### Experimental model

Detailed description of methods employed in this study is provided in SI Appendix.

*Trypanosoma brucei* subsp. *brucei*, strain Lister 427 29-13 (TetR T7RNAP) is a procyclic form cell line that expresses T7 RNA polymerase (T7RNAP) and tetracycline repressor (TetR). Strain Lister 427 29-13 (TetR T7RNAP) was derived by sequential sDataset transfections of the procyclic Lister 427 strain (BEI Resources NR-42010, (38)). This cell line was maintained in SDM-79 media supplemented with neomycin, hygromycin and 10% fetal bovine serum at 27°C. Protein and RNAi expressing transgenic cell lines were maintained in the same media with phleomycin. MERS1-KI and MERS1 mut-KI cell lines were maintained in the same media as TAP and RNAi strains supplemented with blasticidin, puromycin and tetracycline.

### Inducible RNAi and knock-in cell lines

Plasmids for RNAi knockdowns were generated by cloning ~500-bp gene fragments into p2T7-177 vector for tetracycline-inducible expression (39). Linearized constructs were transfected into a procyclic 29-13 *T. brucei* strain (38). For inducible protein expression, full-length genes were cloned into pLew-MHTAP vector (40). Generation of MERS1-KI and MERS1 mut-KI cell lines is described in Appendix.

### Protein purification and analysis

Mitochondrial isolation, glycerol gradient fractionation, native gel electrophoresis, Western blotting, tandem and rapid affinity protein purification, immunoprecipitation and mass spectrometry were performed as described (41).

### RNA purification and analysis

Quantitative RT-PCR, Northern blotting, 3′ RACE, CLIP-Seq and CLAP-Seq have been described previously (19). For 5′ RACE, 5 μg of total RNA was treated with RNA 5′ polyphosphatase (Epicentre) and RNA was ligated with 50 pmol of RA5 RNA 5′ adapter (Illumina) using T4 RNA ligase 1 (New England Biolabs). *In vitro* transcribed luciferase mRNA fragment was used as spike (1 ng), and the equivalent of 2 μg of RNA was used to generate cDNA. Libraries were amplified with Illumina universal forward primer and Illumina indexed reverse primers. For the 5′ RACE analysis of sense maxicircle encoded transcripts, 4 biological replicate experiments were performed with RNA extracted from cells cultured at different times.

### *In vivo* RNA stability assay

The conditions were adapted from (17). RNAi was induced with 1 μg/ml of tetracycline in 100 ml culture (6×10^5^ cells/ml) and cultivation continued for 55 h, with cultures typically reaching cell density of ~5×10^6^/ml. Actinomycin D and ethidium bromide were added to 20 μg/ml and 10 μg/ml, respectively, to block transcription. Cells were collected in 15 ml aliquots after 0.5, 1, 2, and 4 hours by centrifugation at 3000 g for 10 min, washed with ice-cold PBS, re-pelleted and frozen in liquid nitrogen. RNA was isolated and analyzed by Northern blotting in 5% PAGE with 8M urea. The change in relative abundance was calculated assuming the mRNA/tRNA ratio in mock-induced cells at the time of Actinomycin D addition as 100%.

### Kinetoplast affinity purification–sequencing (KAP-Seq)

Live cells were resuspended in 40 ml of SDM79 media at 10^7^/ml and mixed with 4 ml of crosslink solution (50 mM HEPES pH 7.3, 100 mM NaCl, 1 mM EDTA, 2.5% formaldehyde). Suspension was incubated at room temperature for 20 min with mixing. The crosslinking reaction was quenched with 2.5 ml of 2M glycine. Cells were pelleted by centrifugation for 15 min at 3000 g, 4°C, washed with 50 ml of PBS and flash frozen with liquid nitrogen, and processed as described in Appendix.

### *In vitro* activity assays

The following RNA substrates were prepared by *in vitro* transcription:

ND8 mRNA fragment (maxicircle position 3165-3202):

GAAUCAAUUUAAUAAUUUUAAGUUUUGGUUGAUUAAAA
RPS12 and ND5 junction region (maxicircle position 14335-14534):

GGAGAGAAAGAGCCGUUCGAGCCCAGCCGGAACCGACGGAGAGCUUCUUUUGAA
UAAAAGGGAGGCGGGGAGGAGAGUUUCAAAAAGAUUUGGGUGGGGGGAACCCUU
UGUUUUGGUUAAAGAAACAUCGUUUAGAAGAGAUUUUAGAAUAAGAUAUGUUUU
UAAUAUUUUUUUUAUUUUUUAUAAUGUUUGGGUUUAUA
Antisense RNA for RPS12 mRNA (maxicircle position 14418-14335):

GGUUUGAAACUCUCCUCCCCGCCUCCCUUUUAUUCAAAAGAAGCUCUCCGUCGGU
UCCGGCUGGGCUCGAACGGCUCUUUCUCUCC
ND7 and CO3 junction region (maxicircle position 4674-4879):

GGAAUUUUUGGGGGAGCUCGACGGCGGGCGGAGCAUUAUUUGAGGAGGGCGGGA
GCAGAAGGCUUUCUGAGGAAAGAGGGGACCGAGAUCGAUGAAGGUUAUUUUUUG
GUUAUUGAGGAUUGUUUAAAAUUGAAUAAAAAGGCUUUUUGGAAGGGGAUUUUU
GGGGGACACCGCCAGAGGAGGAGGGUUUUGGAAGAGUUUGUUUU
Antisense RNA for ND7 mRNA (maxicircle position 4743-4674):
GGAGAAAGCCUUCUGCUCCCGCCCUCCUCAAAUAAUGCUCCGCCCGCCGUCGAGC
UCCCCCAAAAAUUCC

Triphosphorylated ND8 transcript bearing a 5′-terminal γ-^32^P was synthesized in 100 μl reaction containing 40 mM Tris-HCl pH 7.8, 20 mM NaCl, 26 mM MgCl_2_, 2 mM spermidine, 10 mM DTT, 0.1% Triton X-100, 80 U T7 RNA polymerase (Ambion), 1 μM template, 0.5 mM ATP, 0.5 mM CTP, 0.5 mM UTP, 12.5 μM unlabeled GTP and 0.42 μM [γ-^32^P] GTP (250 μCi, Perkin Elmer).

Pyrophosphohydrolase activity assay was carried out in 20 μl reaction containing 50 mM HEPES pH 7.5, 100 mM NaCl, 2 mM MgCl_2_, 0.5 mM MnCl_2_, 1 mM DTT, 50,000 cpm of ^32^P-γ labeled ND8 substrate, and 4 μl of TAP-purified complexes fractions (MERS2, GRBC5, RGG2, and LSU-4710) or rapid affinity-purified MERS1 fraction. The reaction mixture was pre-incubated at 30°C for 10 min, and the reaction was started by addition of the RNA substrate. To identify the released pyrophosphate and orthophosphate, the same substrate was treated with 0.04 U/μl of 5′ RNA pyrophosphohydrolase (RppH, New England Biolab) in 20 μl reaction containing 10 mM Tris-HCl pH 7.9, 50 mM NaCl, 10 mM MgCl_2_, 1 mM DTT at 30 °C for 20 min. Aliquots (5 μl) were taken in 10, 20 and 30 min, and transferred into 2 μl of 2 μg/μl Proteinase K (Thermo Fisher) and kept on ice until last time point completed. All aliquots were incubated at 30°C for 10 min to digest proteins in the reaction, and 1.5 μl of each aliquot was analyzed by thin-layer chromatography on a PEI-cellulose F plate (Millipore) developed with 0.3 M potassium-phosphate buffer pH 7.4 for 1 hour. TLC plates were air-dried and exposed to phosphor storage screens. Phosphor images were acquired with Typhoon FLA 7000 (GE Healthcare). *In vitro* MPsome activity was tested as described (16).

## Acknowledgments

We thank members of the Aphasizhev laboratory for valuable discussions. We thank Ken Stuart for providing pSM06 and pSM07 plasmids. This work was supported by NIH grants RO1 AI113157 to IA, RO1 GM074830 and RO1 GM106003 to LH, and RO1 AI091914 and RO1 AI101057 to RA.

## Accession Numbers

RNA-Seq and KAP-Seq data files were deposited to BioProject (http://www.ncbi.nlm.nih.gov/bioproject/) under accession number PRJNA402081.

## Author Contributions

FS conducted KAP-Seq, CLIP, RACE and characterized native complexes. TS carried out *in vitro* reconstitutions. LZ and TY performed bioinformatics analysis. LH directed protein identification by mass spectrometry. IA performed genetic knockdowns and affinity purifications. FS and RA wrote the manuscript. All authors contributed to the final version of the paper.

## Conflict of Interest

The authors declare that they have no conflict of interest.

---

**Figure S1.**
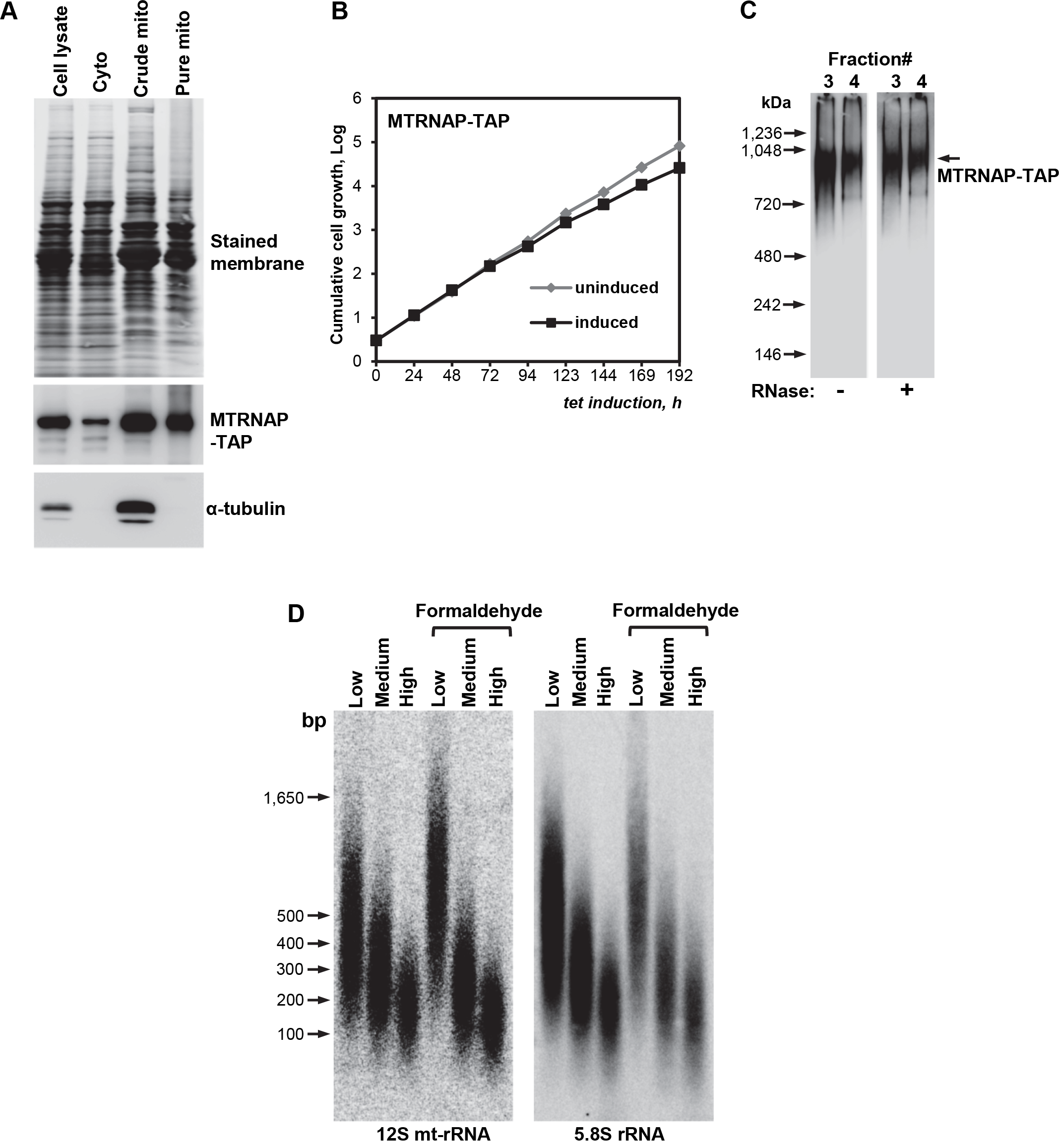
MTRNAP Complex Characterization. Related to Figure 1. (A) Mitochondrial localization of MTRNAP-TAP. Subcellular fractionation was performed with MTRNAP-TAP expressing cell line. Cyto, cell lysate depleted of membranes by centrifugation; crude mito, mitochondria and cell membranes; pure mito, mitochondrial fraction purified by centrifugation in Renografin density gradient. Protein profiles were visualized by Sypro Ruby staining and specific proteins were detected by immunoblotting. (B) Growth kinetics of MTRNAP-TAP cell line after induction with tetracycline. (C) Native molecular mass of the MTRNAP complex. Mock-and RNase I-treated cell lysates were separated on a 10 – 30% glycerol gradients. Peak fractions were resolved on native gel alongside with molecular mass markers. MTRNAP-TAP was detected by immunoblotting. The fraction 3,4 are the same fractions from Figure 1A. (D) Southern blotting analysis of maxicircle DNA fragments generated by focused sonication in mock-and formaldehyde-crosslinked cells. Low, medium and high intensity shearing was performed with Covaris M220 Focused Ultrasonicator as described in Star*Methods. High intensity was used to generate KAP-Seq libraries.

**Figure S2.**
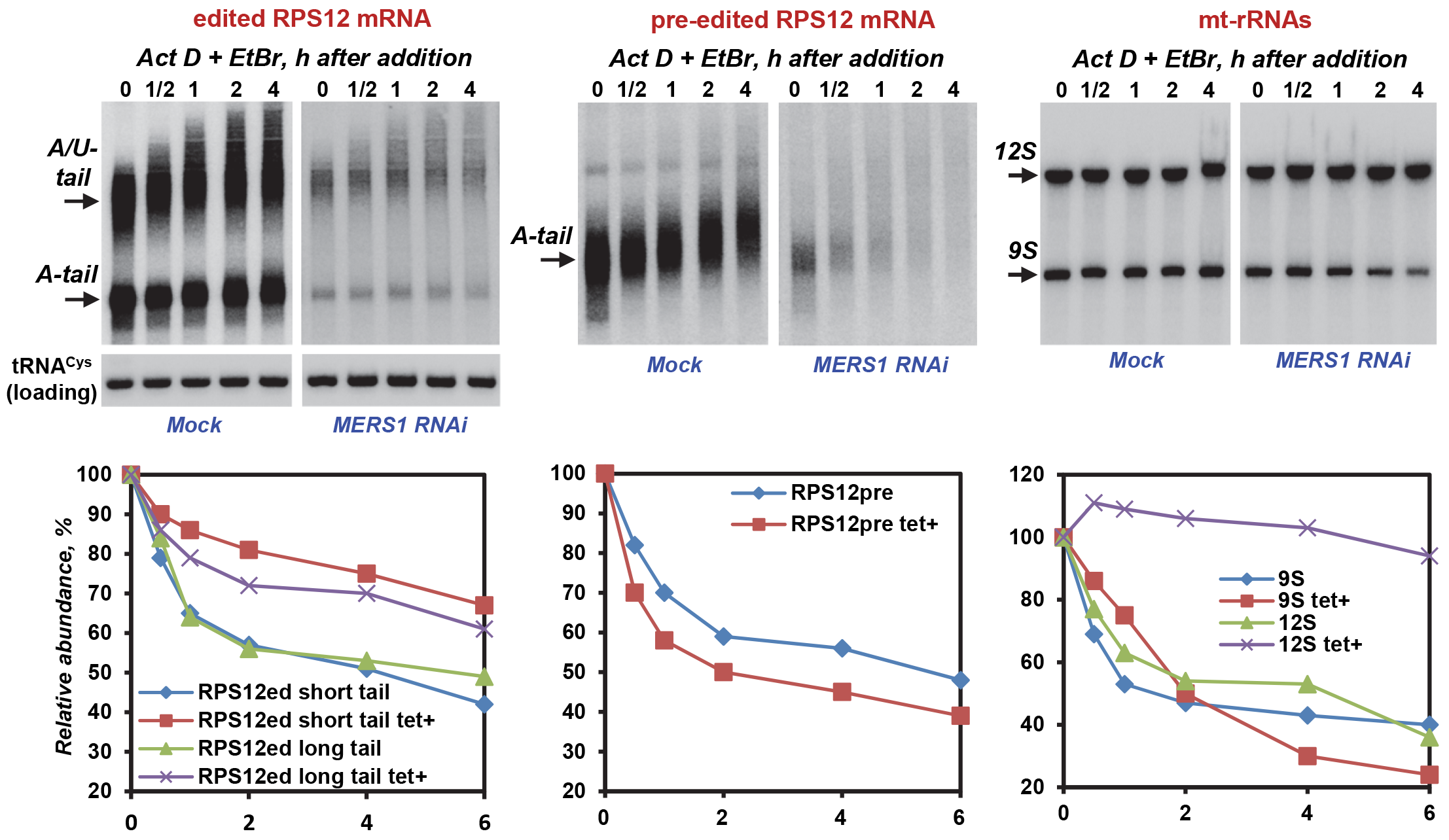
Real-time RNA decay in MERS1 RNAi cells. Related to Figure 4. After depleting MERS1 by inducible RNAi for 55 hours, Actinomycin D and ethidium bromide were added to inhibit transcription (zero-time point). Total RNA was isolated at indicated time intervals after ActD/EtBr bromide addition, separated on denaturing PAGE and sequentially hybridized with DNA probes for fully-edited and pre-edited RPS12 mRNAs, and 9S and 12S mt-rRNAs. Nuclear-encoded tRNA^Cys^ that is predominantly localized in the mitochondrion was used as loading control. The graphs corresponding to Northern blotting panels represent changes in relative abundance, assuming the mRNA/tRNA ratio at the time of ActD/EtBr addition as 100%. Note pre-edited mRNA lengthening in mock-induced cells over the assay duration; this indicates unperturbed U-insertion editing under transcription blockade conditions.

**Figure S3.**
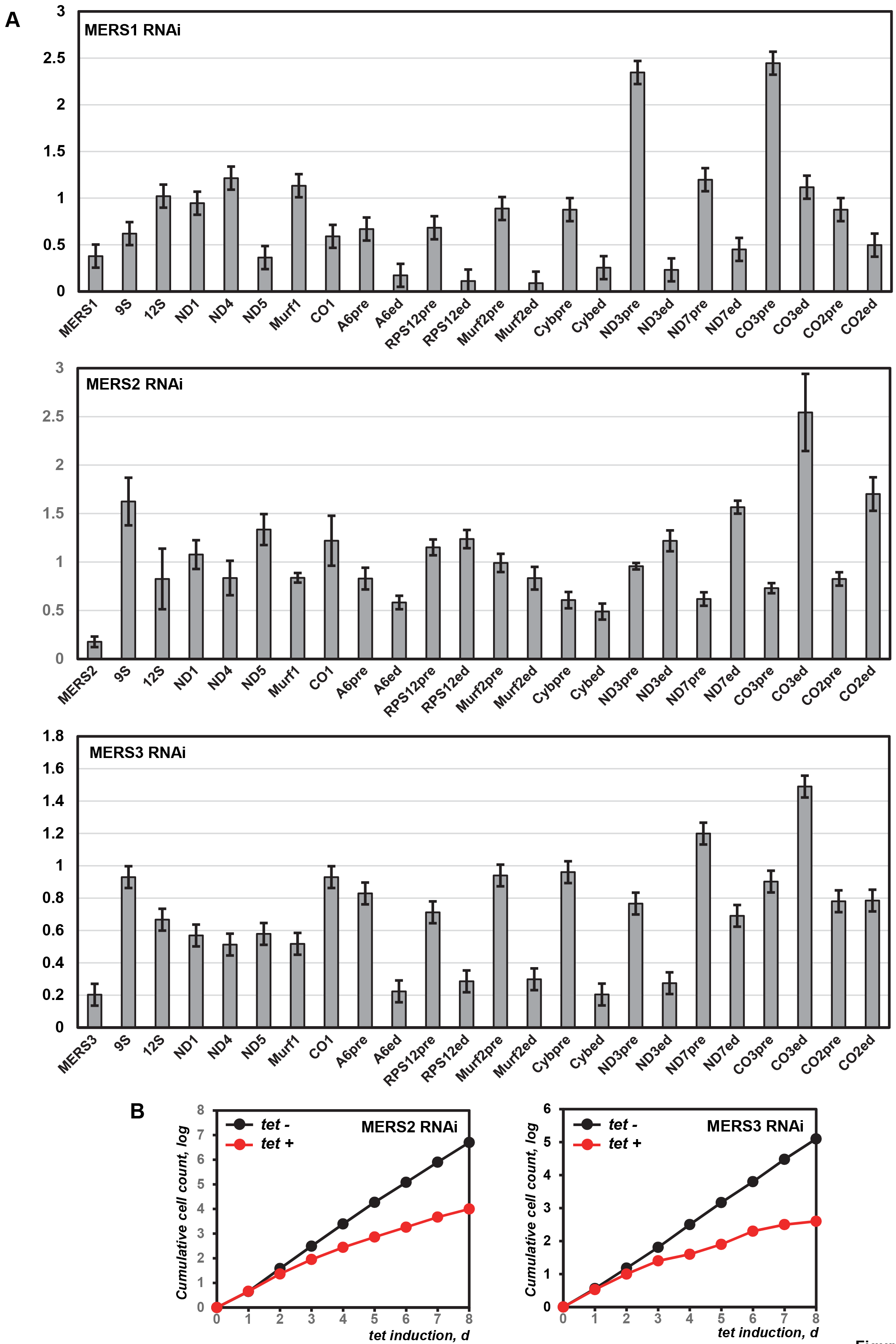
Effect of Individual PPsome Subunit Knockdowns on Maxicircle-Encoded Transcripts and Cell Growth. Related to Figure 4. (A) Quantitative real-time RT-PCR analysis of RNAi-targeted MERS1, MERS2 and MERS3 mRNAs, and mitochondrial rRNAs and mRNAs. The assay distinguishes edited and corresponding pre-edited transcripts, and unedited mRNAs. RNA levels were normalized to β-tubulin mRNA. RNAi was induced for 72 hours. Results are presented as mean of three technical replicates ± SD. Line at “1” reflects no change in relative abundance; bars above or below represent an increase or decrease, respectively. Pre, pre-edited mRNA; ed, edited mRNA. (B) Growth kinetics of parasite suspension cultures after RNAi induction.

**Figure S4.**
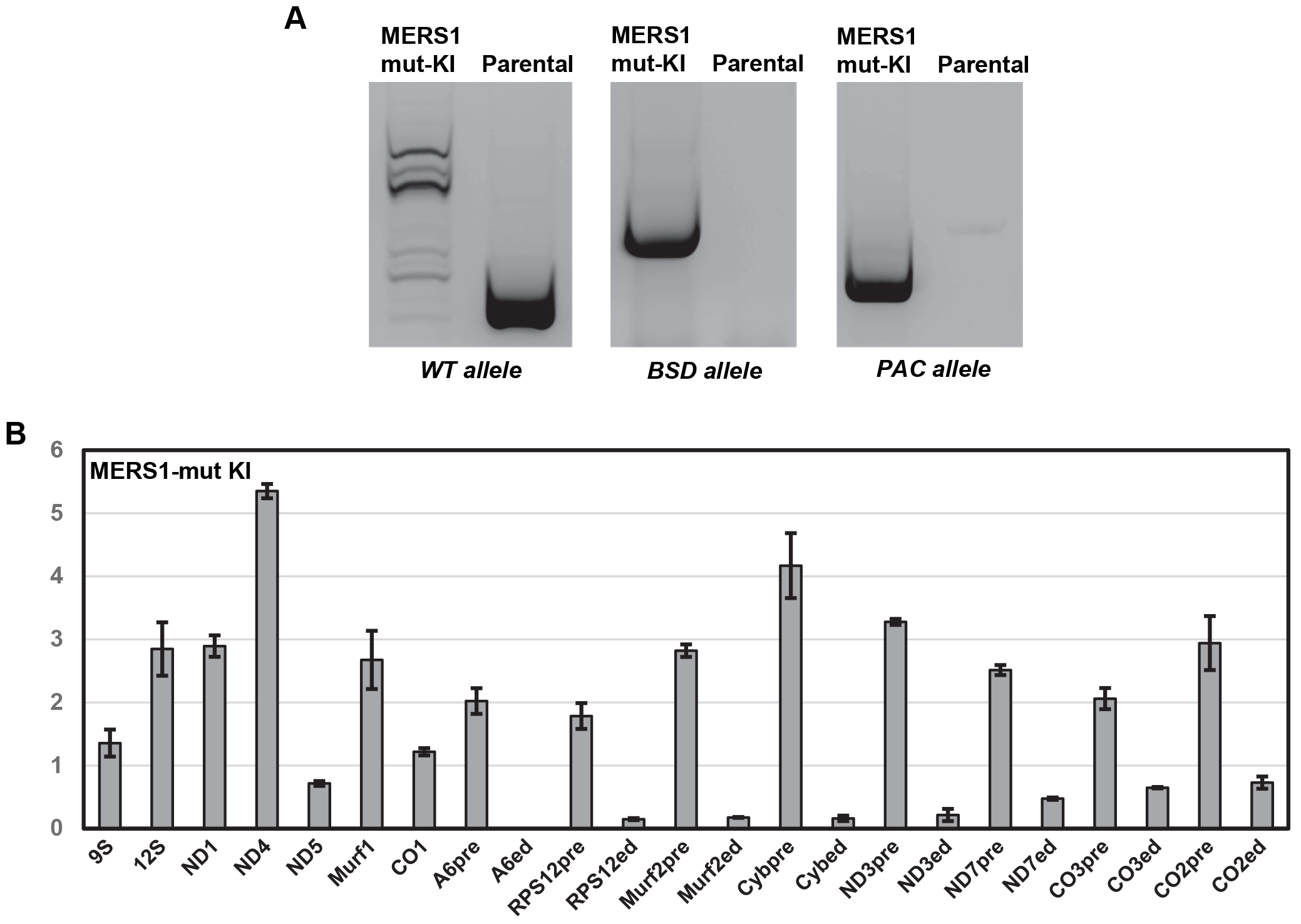
Characterization of MERS1 knock-in cell lines. Related for Figure 5. (A) Disruption of the first allele by Blasticidin S (BSD) resistance gene, and replacement of the second allele with a cassette constitutively expressing mutated MERS1 (puromycin resistance, PAC). Genomic PCR amplification products were separated on agarose gel. (B) Quantitative real-time RT-PCR analysis of mitochondrial rRNAs and mRNAs in MERS1-mutKI cell line. The assay distinguishes edited and corresponding pre-edited transcripts, and unedited mRNAs. RNA levels were normalized to β-tubulin mRNA. The tetracycline was withdrawn for 120 hours. Results are presented as mean of three technical replicates ± SD. Line at “1” reflects no change in relative abundance; bars above or below represent an increase or decrease, respectively. Pre, pre-edited mRNA; ed, edited mRNA. The edited A6 mRNA was downregulated beyond detection level.

## Supplemental Information

### Detailed protocols

#### Inducible RNAi, protein expression and knock-in cell lines

Plasmids for RNAi knockdowns were generated by cloning ~500-bp gene fragments into p2T7-177 vector for tetracycline-inducible expression (Wickstead et al., 2002). Linearized constructs were transfected into a procyclic 29-13 *T. brucei* strain (Wirtz et al., 1999). For inducible protein expression, full-length genes were cloned into pLew-MHTAP vector (Jensen et al., 2007). Expression was induced by adding tetracycline to 1 mg/L.

To generate the MERS1-KI and MERS1 mut-KI cell lines, two constructs were sequentially transfected into cell line that expresses MERS1-TAP in pLew-MHTAP background. The first construct, MERS1BSD, contained MERS1 5′ UTR, the blasticidin resistance gene and MERS1 3′ UTR. Each DNA fragment was independently amplified from genomic total DNA or pSM06 plasmid (Blasticidin resistance, (Merritt and Stuart, 2013)) and gel-purified. To generate MERS1BSD, 10 ng of each PCR product was mixed in the same PCR reaction and the final construct was amplified using forward primer and reverse primer used to amplify MERS1 5′ UTR and 3′ UTR, respectively. After gel purification, the PCR product was cloned into pCR2.1 TOPO vector (Thermo Fisher, Cat# 451641). Clonal cell lines (MERS1 TAP MERS1BSD) were obtained by limiting dilution in the presence of blasticidin and tetracycline. Knockout of the first allele was assessed by PCR. The second construct contained MERS1 5′UTR, MERS1 or MERS1 mut gene (E257A, E258A), tubulin spacer region, puromycin resistance gene and MERS1 3′UTR in a pUC19 vector backbone. Individual fragments were amplified from genomic DNA or pSM07 plasmid (puromycin resistance gene, (Merritt and Stuart, 2013)) and used for Gibson Assembly. The MERS1 PAC (with MERS1 gene) and MERS1 mut PAC (with MERS1 mut gene, E257A, E258A) constructs were transfected in MERS1TAP MERS1 BSD cell line in the presence of tetracycline. After clonal selection, the loss of wild type MERS1 alleles and integration of each construct in the MERS1 loci was assessed by PCR.

#### Western blotting

Proteins were separated on precast 8%-16% SDS Tris-glycine polyacrylamide gels (Thermo Fisher) and transferred onto nitrocellulose membrane. Detection was performed with rabbit antigen-purified rabbit polyclonal antibodies (MERS1, KPAP1 and GRBC1-2), mouse monoclonal antibody (RET1), or Peroxidase Anti-Peroxidase Soluble Complex antibody to detect TAP tagged proteins. Quantitative chemiluminescent images were acquired with LAS-4000 imager (GE Healthcare).

#### RNA isolation

Cell culture equivalent of 25×10^7^ cells was collected by centrifugation at 3000 g for 10 min at 4°C, washed with 50 ml of PBS, collected likewise, transferred to 2 ml tube, and re-pelleted at 3000 g for 5 min at 4°C. Cell pellets were resuspended in 0.8 ml of cold Solution D (4M guanidine isothiocyanate, 25 mM sodium citrate, pH 7.0, 0.5% sarcosyl, 0.1 M 2-mercaptoethanol) and supplemented with 0.1 ml of 2 M sodium acetate (pH 4.0) and 0.9 ml of water-saturated phenol, and gently mixed. After addition of 0.3 ml of chloroform/isoamyl alcohol (49:1), lysates were incubated for 10 min on Nutator at 4°C. Phases were separated by centrifugation at 21,000 x g for 10 min, and the supernatant was transferred into a Phase Lock Gel Heavy 2-ml tube (5Prime) and extracted vigorously by vortexing for 1 min with 0.8 ml of chloroform/isoamyl alcohol. Supernatant was transferred into a fresh tube and RNA was precipitated with 1 ml of isopropanol at −20°C for 1 hour. RNA was collected by centrifugation at 21,000 x g for 15 min and washed with 80% ethanol. RNA was dissolved in 0.2 ml of water and re-precipitated with sodium acetate/ethanol by standard technique. For Northern blotting, 50 μg was treated with 5U of RNase-free DNase I (New England Biolabs) in 0.1 ml of manufacturer-supplied buffer for 30 min at 37°C. RNA was extracted with equal volume of phenol (pH 5), and phenol-chloroform, and precipitated with ethanol. For quantitative RT-PCR, a Turbo DNase digestion was performed for 30 min at 37°C and RNA was purified with RNeasy MiniElute Kit (Qiagen) as recommended by the manufacturer.

#### Northern blotting

Northern blotting experiments were performed according to published protocol (Aphasizheva et al., 2016). Total RNA (8 ‑ 12 μg) was separated on 20 cm-long 5% (mRNA and rRNA detection) or 10% (tRNA and gRNA detection) acrylamide/ bis-acrylamide (19:1)/8M urea gel. RNAs were transferred to a BrightStar®-Plus membrane (Thermo Fisher Cat# AM10104) by tank transfer at 100 V for 2 hours at 4°C. Alternatively, total RNA was separated on 20 cm-long 1.7% agarose/formaldehyde gels (CO1 and cyb mRNA detection). After electrophoresis, RNA was transferred to a BrightStar®-Plus membrane (Thermo Fisher Cat# AM10104) using vacuum blotting system at 50 mbar for at least 4 hours or overnight. After transfer, RNA was cross-linked to membrane by exposition to UV light at 120 mJ/cm^2^ using CX-2000 UV Crosslinker (UVP).

Oligo probes were labeled by incubating 10 pmol of oligonucleotide with T4 polynucleotide kinase (Ambion) in the presence of 6.5 μl of [γ-32P] ATP (6000 μCi/ml) (Perkin Elmer) for 30 min at 37°C. Labeled oligoprobe was purified on G-25 Sephadex column (GE Life Sciences). For single-stranded DNA probe, the template was amplified with oligos used for quantitative RT-PCR. DNA recovered from a single 25 μl qRT-PCR reaction was typically sufficient for 20 single-strand probe preparations by asymmetric two-step PCR: 10 pmol of 5-radiolabeled antisense oligonucleotide in a standard PCR reaction with Taq polymerase (95°C, 60 s; 95°C, 15 s; 50°C, 30 s; 45 cycles). The probe was purified on G25 Sephadex spin column (GE Life Sciences). For the detection of antisense mRNA, the radiolabeled ssDNA probe was produced using the sense oligonucleotide (instead of antisense) in the PCR reaction.

Hybridization with radiolabeled probes was performed overnight using ULTRAhyb ™ Ultrasensitive Hybridization buffer (ThermoFisher Cat# AM8670) or ULTRAhyb ™-Oligo Hybridization buffer (ThermoFisher Cat# AM8663). Oligonucleotides and ssDNA PCR products were hybridized overnight at 40°C and 42°C, respectively. Membranes were washed 3 times at the hybridization temperature with 15 ml of 2x SSPE 0.5% SDS or 4x SSPE 0.5% SDS for PCR probes and oligonucleotides, respectively. Membranes were exposed to a phosphor storage screen.

#### In vivo RNA stability assay

The conditions were adapted from (Aphasizheva and Aphasizhev, 2010). RNAi cells were stabilized in SDM79 media with 10% serum, 50 μg/ml of hygromycin, 30 μg/ml of G418 and 2.5 μg/ml of phleomycin. RNAi was induced with 1 μg/ml of tetracycline in 100 ml culture (6×10^5^ cells/ml) and cultivation continued for 48 h, with cultures typically reaching cell density of ~5×10^6^/ml. Actinomycin D and ethidium bromide were added to 20 μg/ml and 10 μg/ml, respectively, to block transcription. Cells were collected in 15 ml aliquots after 30 min, 1, 2, and 4 hours by centrifugation at 3000 μg for 10 min, washed with ice-cold PBS, re-pelleted and frozen in liquid nitrogen. Total RNA was isolated and analyzed by Northern blotting in 5% PAGE with 8M urea. The change in relative abundance was calculated assuming the mRNA/tRNA ratio in mock-induced cells at the time of Actinomycin D addition as 100%.

#### 5′ RACE

After a DNase treatment (as for qRT-PCR, RNA isolation protocol), 5 μg of total RNA was treated with RNA 5′ polyphosphatase (diP treatment, Epicentre) according to manufacturer recommendations, extracted by phenol/chloroform and precipitation with ethanol. RNA was ligated with 50 pmol of RA5 RNA 5′ adapter using T4 RNA ligase 1 (New England Biolabs) in 50 μl reaction for 16 hours at 16°C. Spike RNA (*in vitro* transcribed Luciferase mRNA fragment) was added prior to ligation (10 ng for sense mRNA 5′end detection, and 0.5 ng for antisense mRNA detection). After phenol/chloroform extraction and ethanol precipitation, the equivalent of 2 μg of RNA was used to generate cDNA with 2 pmol of each gene-specific primer with 200 U of Superscript III (Thermo Fisher). The same reactions were performed in the absence of Superscript III (−RT controls). After cDNA synthesis, 0.5 U of RNase H (Thermo Fisher) was added and incubation continued for 30 min at 37°C. Libraries were amplified in a 50 μl PCR reaction in with 600 pmol of each Illumina universal forward primer and Illumina indexed reverse primers, using 2.5 μl of cDNA and Phusion Hot Start DNA polymerase II (Thermo Fisher) supplemented with1 μg of RNase A (Qiagen). After an initial denaturation step of 30 s at 98°C, the PCR reactions underwent 13 PCR cycles (98°C 15 s, 60°C 30 s, 72°C 15 s) and a final elongation step of 30 s at 72°C. PCR products were purified on a ZYMO DNA clean up and concentrator-5 column, eluted in 10 μl, and resolved on a 7.5% acrylamide/TBE gel. After staining the gel with SYBR Green 1, the area of interest was excised under blue light and PCR products were eluted, precipitated and purified with ZYMO DNA clean up and concentrator-5 column. For the 5′ RACE analysis of sense maxicircle encoded transcripts, 4 biological replicate experiments were performed with RNA extracted from cells cultured at different times and started from different frozen aliquots.

#### 3′ RACE

Total RNA (5 μg) was treated with Calf Intestinal Alkaline Phosphatase (Thermo Fisher) according to manufacturer recommendations. After phenol/chloroform extraction and ethanol precipitation, RNA was ligated to 100 pmol of RA3 RNA 3′ adapter with T4 RNA ligase 1 (New England Biolabs) in a 50 μl reaction, overnight at 16°C. Spike RNA was added prior to ligation (10 ng). After phenol/chloroform extraction and ethanol precipitation, 2 μg was used to generate cDNA using the RTP primer and 200U of Superscript III (Thermo Fisher). After cDNA synthesis, 0.5 U of RNase H (Thermo Fisher) was added to the cDNA synthesis reaction and incubated for 30 min at 37°C. First PCR amplification was performed in 100 μl reaction with 2 μl of cDNA as template, 600 pmol of Illumina indexed reverse primer and 20 pmol of each gene-specific primer, 1 μg of RNase A (Qiagen), using Phusion Hot Start II DNA polymerase. After an initial denaturation step of 30 s at 98°C, PCR reactions underwent 5 PCR cycles at a low hybridization temperature (98°C 15 s, 50°C 30 s, 72°C 15 s) and 5 PCR cycles at a high hybridization temperature (98°C 15s, 60°C 30 s, 72°C 15 s) and a final elongation step at 72°C for 30 s. After PCR products purification on a ZYMO DNA clean and concentrator-5 column and elution in 20 μl, libraries were amplified in a 50 μl PCR reaction in the presence of 600 pmol of each Illumina universal forward primer and Illumina indexed reverse primers, using 2 μl of the first PCR as template and Phusion Hot Start DNA polymerase II (Thermo Fisher). After an initial denaturation step of 30 s at 98°C, the PCR reactions underwent 16 PCR cycles (98°C 15 s, 60°C 30 s, 72°C 15 s) and a final elongation step of 30 s at 72°C. After purification on a ZYMO DNA clean and concentrator-5 column, the PCR products were resolved on a 7.5% acrylamide/TBE gel. After staining the gel with SYBR Green 1, the area of interest was excised under blue light and eluted, precipitated and purified with ZYMO DNA clean up and concentrator-5 column.

#### Quantitative RT-PCR

cDNA was synthesized from 2 μg of Turbo DNase-treated, column purified total RNA in 0.1 ml reaction with TagMan Reverse Transcription Reagents (N808-0234, Applied Biosystems) as recommended by the manufacturer. Primer pairs were pre-mixed at 1.5 μM final concentration of each in water. In a 1.5 ml tube, 8 μl of cDNA was mixed with 18 μl of primer mix. Power SYBR Green Master Mix was added to 50 μl. Triplicate aliquots of 16 μl were distributed into 96 well plate (951022043, Eppendorf). After sealing the plate with a film (951023060, Eppendorf), PCR reactions were performed in Eppendorf Realplex 2S cycler as follows: 95°C, 10 min; 95°C (15 s), 60°C (1 min, measure point), 45 cycles.

#### Mitochondrial preparations

Crude fractionation. Parasite culture was inoculated at 10^6^ cells/ml and grown in 800 ml of SDM-79 media with 10% FBS and required antibiotics at 27°C in a roller bottle at 8 rpm to 15-20×10^6^ cells/ml (60-72 hours). Cells were collected by centrifugation at 3000g for 10 min at 4°C. Cell pellet was resuspended and washed in 50 ml of phosphate buffered saline (PBS) plus 6 mM sucrose. Cells were resuspended in DTE buffer (5 mM Tris-HCl, 1mM EDTA, pH 8) to achieve final concentration of 1.2×10^9^ cells/ml. During the centrifugation steps, a 50 ml conical tube (rated at g-force of 15,000 or higher) was prepared with pre-calculated volumes of 60% sucrose (12 ml of sucrose per 100 ml of DTE), 150 μl of 1M MgCl_2_, and 0.2 ml of DNase I solution (5000 U/ml, Sigma, Cat# D5025). Cell suspension in DTE was transferred into 10 ml syringe fitted with 26-gauge needle and push intensely into the prepared sucrose cushion. After gentle mixing, the total volume was brought to 50 ml with STE buffer (20 mM Tris-HCl pH 7.6, 250 mM sucrose, 1 mM EDTA) and lysate was incubated on ice for 15 min. The crude mitochondria pellet was collected by centrifugation at 15,000g for 15 min at 4°C. After a wash step with 50 ml of STE, crude mitochondria pellet was flash-frozen in liquid nitrogen, or used for pure mitochondria preparation.

Density gradient mitochondria preparation. Crude mitochondria pellet was resuspended in 2 ml of 76% RSTE (76% Renografin (Bracco Diagnostics, Ren°Cal 76 Cat# 086032)) in STE buffer, and loaded at the bottom of a 20 – 35% Renografin gradient formed in 20 mM Tris-HCl pH 7.9, 250 mM sucrose, 1 mM EDTA (SW-28 rotor tubes, Beckman). After centrifugation for 2 hours at 24,000 rpm, 4°C, the band corresponding to pure mitochondria (usually in the middle of the gradient) was harvested by side puncture with 18# needle and transferred in to 50 ml tube. Mitochondrial fraction was diluted 5-fold with cold STE buffer and centrifuged for 20 min at 15,000g, 4°C. After a final wash with 2 ml of STE, pure mitochondria were centrifuged again for 20 mins at 15,000g, 4°C before flash freezing the pellet with liquid nitrogen.

#### Glycerol gradient fractionation and immunoprecipitation

Cell pellet (~5×10^8^ cells), or 200 mg (wet weight) of crude mitochondrial pellet, was resuspended in 0.3 ml of Gradient Lysis Buffer (GLB, 30 mM HEPES, pH 7.3, 120 mM KCl, 12 mM MgCl_2_, 1 mM DTT, 1/10 of Complete Protease Inhibitor, 2 U of Turbo DNase and 1.2% NP40) and incubated on ice for 10 min. The lysate was centrifuged for 15 min at 21,000g and the supernatant was recovered. For RNase treatment, 100 U of RNase I was added to the lysate and incubated on ice for 10 min. The10%-30% glycerol gradient in 25 mM HEPES pH 7.3, 100 mM KCl and 10 mM MgCl_2_ was prepared for SW41 rotor (Beckman) tubes using Gradient Master (Biocomp Instruments). The extract (0.25 ml) was centrifuged for 5 hours at 38,000 rpm in SW41 rotor (Beckman) with slow breaking and 0.56 ml fractions were collected from the top with Gradient Station fractionator (Biocomp Instruments). Each fraction (10 μl) was supplemented with Coomassie R250 to 0.25%, separated on precast NativePAGE 3%-12% Bis-Tris Protein Gels (Thermo Fisher) and transferred onto nitrocellulose membrane for immunodetection.

Immunoprecipitation in glycerol gradient fractions. After mitochondrial lysate fractionation on glycerol gradient, 300 μl from fractions 2-16 were mixed with 200 μl of buffer A (25 mM HEPES pH7.3, 100 mM KCl, 10 mM MgCl_2_, 1.25 mg/ml BSA, 1/100 Complete EDTA-free protease inhibitor). The protein of interest was immunoprecipitated using 0.3 mg of polyclonal antibody-coated Dynabeads (MERS1 IP) or IgG coated magnetic beads (KPAP1-TAP IP) for 1 hour at 4°C on a Nutator platform. Proteins were eluted from beads with 40 μl of 1x SDS gel loading buffer at 70°C for 10 min in a Thermomixer, and separated on precast 8%-16% SDS polyacrylamide gel.

#### Tandem affinity purification

Crude mitochondrial pellet (~1 g wet weight) was resuspended in 3 ml of Lysis Buffer (LB, 50 mM Tris-HCl, pH 7.6, 120 mM KCl, 1% NP40, 5 mM MgCl_2_, 5% glycerol, 1/5 of Complete protease inhibitor tablet (Roche), 20 U Turbo DNase (Thermo Fisher)), and incubated on ice for 15 min. After adding extraction buffer without detergent to 11 ml, mitochondria lysate was sonicated 3 times for 10s at 9W. Extracts were then centrifuged at 200,000g for 15 min in SW41 rotor (Beckman). Supernatant was passed through the low-protein binding 0.45 μm filter and separated in two parts: one was supplemented with 0.1 mg of RNase A and 2000 U of RNase T1, and the other used as control (No RNase). After incubation on ice for 10 min, extracts were transferred to tubes containing 0.2 ml of IgG Sepharose slurry (GE Life Sciences, pre-washed twice with 10 ml of IgG Binding Buffer (IgG-BB, 20 mM Tris-HCl, pH 7.6, 100 mM KCl, 5 mM MgCl_2_, 5% glycerol, 0.1% NP40), and incubated for 30 min at 4°C. The suspension was transferred into 2 ml disposable column with bottom filter and washed with 5 full column volumes (CV) of IgG-BB. After extra wash with 2 CV of IgG-BB plus 1 mM DTT, the column was closed and contents incubated with 150 U of AcTEV protease and 1/200 of Complete inhibitor tablet in 1.5 ml of IgG-BB plus 1 mM DTT. After closing the upper end with Parafilm, the column was incubated for 16 hours at 4 °C on a Nutator platform. The column was drained into 15 ml plastic tube containing 0.2 ml of pre-washed calmodulin resin with calmodulin binding buffer (CBB, 20 mM Tris-HCl, pH 7.6, 100 mM KCl, 2 mM CaCl2, 1 mM MgAc, 0.1% NP40, 10 mM 2-mercaptoethanol, 1 mM imidozole, 5% glycerol). The IgG column was rinsed 4 times with 1 ml of CBB, CaCl2 was added to 1 mM and the suspension was incubated for 1 hour on Nutator at 4°C. The suspension was transferred into 2-ml disposable column and washed with 3 full CV of CBB and protein was eluted with 0.6 ml of Calmodulin Elution Buffer (CEB, 20 mM Tris-HCl, pH 7.6, 100 mM KCl, 2 mM EDTA, 3 mM EGTA, 0.1% NP-40, 10 mM 2-mercaptoethanol, 1 mM imidozole, 5% glycerol).

#### Rapid affinity purification

Procyclic *T. brucei* cells were grown in 850 ml to ~20-25 x 10^6^ cells/ml and harvested by centrifugation at 4,000g for 15 min. Cell pellet was washed with PBS-6 mM sucrose, frozen in liquid nitrogen, powdered with CryoMill (Retsch) and stored at −80°C. All procedures were done at 4°C. All solutions were prepared as described above. Extract was prepared by resuspending frozen powder in 3 ml of pre-warmed lysis buffer (50 mM Tris-HCl pH7.6, 60 mM KCl, 12 mM MgCl_2_, 1% NP-40, 5% glycerol) containing 0.3 ml of Compete protease inhibitor solution and incubated ice for 10 min with 20 U of TURBO DNase (Ambion). Extract was diluted with Extraction buffer (50 mM Tris-HCl (pH7.6), 60 mM KCl, 12 mM MgCl_2_, 5% glycerol) up to 14 ml and centrifuged at 200,000 x g for 30 min. Supernatant was filtered through 0.45 μm low protein binding filter into 15 ml conical tube with 10 mg of IgG-coated Dynabeads magnetic beads (see CLAP protocol). The extract was incubated on Nutator for 20 min. Beads were collected on DynaMag-15 magnet stand (Life Technologies), rinsed twice with 6 ml of wash buffer (20 mM Tris-HCl (pH 7.6), 60 mM KCl, 12 mM MgCl_2_, 0.1% NP-40, 5% glycerol) and washed 3 times with 10 ml of wash buffer for 5 min. Beads were transferred into 1.5 ml low protein binding tube and washed twice with 1 ml of wash buffer using the magnetic holder (Life Technologies). To elute TAP-tagged proteins, beads were incubated with 0.3 ml of TEV digestion buffer (20 mM Tris −HCl (pH 8.0), 0.5 mM EDTA, 1 mM dithiothreitol (DTT)) containing 20 U of TEV protease and 6 μl of Compete protease inhibitor solution for overnight on Nutator at 4°C. Beads were pelleted using the magnetic holder and supernatant was collected as eluate.

#### Mass spectrometry

Affinity-purified complexes were precipitated by addition of trichloroacetic acid and deoxycholate to 20% and 0.1%, respectively, washed three times with ice-cold acetone, and digested with LysC peptidase in 8M urea (1:50 ratio) for 4 hours at 37 °C. Reaction was diluted five-fold with 50 mM Na-bicarbonate pH 7.5 and further digested with trypsin (1:100 ratio) for 16 hours. Peptides were purified on Vivapure spin columns (Sartorius). LC-MS/MS was carried out by nanoflow reversed phase liquid chromatography (RPLC) (Eksigent, CA) coupled on-line to a Linear Ion Trap (LTQ)-Orbitrap mass spectrometer (Thermo-Electron Corp). The LC analysis was performed using a capillary column (100 μm ID x 150 mm) with Polaris C18-A resin (Varian Inc., CA). The peptides were eluted using a linear gradient of 2% to 35% B in 85 min at a flow of 300 nL/min (solvent A: 100% H2O, 0.1% formic acid; solvent B: 100 % acetonitrile, 0.1% formic acid). A cycle of full FT scan mass spectrum (m/z 350-1800, resolution of 60,000 at m/z 400) followed by ten data-dependent MS/MS spectra acquired in the linear ion trap with normalized collision energy (setting of 35%). Target ions already selected for MS/MS were dynamically excluded for 30 s.

Monoisotopic masses of parent ions and corresponding fragment ions, parent ion charge states and ion intensities from the tandem mass spectra (MS/MS) were obtained by using in-house software with Raw_Extract script from Xcalibur v2.4. Following automated data extraction, resultant peak lists for each LC-MS/MS experiment were submitted to the development version of Protein Prospector (UCSF) for database searching similarly as described (Fang et al., 2012). Each project was searched against a normal form concatenated with the random form of the *T. brucei* database (www.genedb.org, v5). Trypsin was set as the enzyme with a maximum of two missed cleavage sites. The mass tolerance for parent ion was set as ± 20 ppm, whereas ± 0.6 Da tolerance was chosen for the fragment ions. Chemical modifications such as protein N-terminal acetylation, methionine oxidation, N-terminal pyroglutamine, and deamidation of asparagine were selected as variable modifications during database search. The Search Compare program in Protein Prospector was used for summarization, validation and comparison of results. Protein identification is based on at least three unique peptides with expectation value ≤ 0.05.

### Coupled Transcription/Translation System and immunoprecipitation

TNT reactions were set up as suggested by manufacturer’s protocol (Promega) with 0.2 μg of each plasmid and [^35^S]-Methionine in 10 μl reactions. Protein G Dynabeads (Life Technologies) were pre-washed with Immunoprecipitation buffer containing 20 mM Tris-HCl (pH 7.5), 150 mM NaCl, 0.2% Tween and 0.5 mg/ml BSA, and resuspended in Immunoprecipitation buffer at 30 mg/ml. All incubations of Dynabeads were performed by using Thermomixer (Eppendorf) at 1000 RPM. After the reaction mixtures were incubated for 90 min at 30 °C, 150 μg of the pre-washed beads was added to each reaction and incubated for 30 min at 250C in order to minimize non-specific binding proteins to the beads. Beads were separated by a magnetic holder, and the supernatant was transferred to a new tube. To analyze the product, 2 μl of the supernatant was mixed with 18 μl of SDS loading buffer (Life Technologies) and heated at 65 °C for 20 min. The remaining supernatant was incubated with 2 μg of MERS1 polyclonal antibody at 25 °C for 30 min. After the incubation with the antibody, 300 μg of the pre-washed beads was added and further incubated at 25 ° C for 10 min. Beads were washed 5 times with Immunoprecipitation Buffer and then resuspended in 20 μl of Immunoprecipitation buffer. To analyze bound proteins, 5 μl of the resuspension was mixed with 15 μl of SDS loading buffer and heated at 65 °C for 20 min. Samples were separated on 8-16% Tris-Glycine gel, transferred on Nitrocellulose membrane (Bio-Rad) and exposed to phosphor storage screen.

#### Kinetoplast Affinity Purification–Sequencing (KAP-Seq)

Live cells were resuspended in 40 ml of SDM79 media at 10^7^/ml and mixed with 4 ml of crosslink solution (50 mM HEPES pH 7.3, 100 mM NaCl, 1 mM EDTA, 2.5% formaldehyde). Suspension was incubated at room temperature for 20 min with mixing. The crosslinking reaction was quenched with 2.5 ml of 2M glycine. Cells were pelleted by centrifugation for 15 min at 3000 g, 4°C, washed with 50 ml of PBS and flash frozen with liquid nitrogen. Pellets were resuspended in 300 μl of Lysis Buffer (LB: 50 mM Tris pH 7.5, 300 mM NaCl, 5 mM EDTA, 1% NP40, 0.1% sodium deoxycholate) and sonicated with Covaris M220 Focused Ultrasonicator in a microTUBE AFA Fiber Screw-Cap (Covaris Cat# 520096) with the following settings: Peak incident power 50 W, duty factor 40%, cycle/burst 200/450 seconds (high, Fig S1D)). For the other sonication conditions presented in Fig S1D, low sonication conditions were: Peak incident power 75W, duty factor 5%, cycle/burst 200/120 seconds. Medium sonication conditions were: Peak incident power 75W, duty factor 20%, cycle/burst 200/240 seconds. The “high” sonication condition was selected for further experiments. After sonication, the lysate was diluted with 5 volumes of LB without detergent (50 mM Tris pH 7.5, 300 mM NaCl, 1 mM EDTA) and centrifuged at 21,000 G for 30 min at 4°C. The supernatant was mixed with 3 mg of Dynabeads™ Protein G cross-linked to Anti-Protein A antibody (Sigma Cat# P3775) using disuccinimidyl suberate (DSS). After adding 1 μg of RNase A, the mixture was incubated overnight at 4°C on a Nutator platform. Beads were the washed 4 times with Wash Buffer 1 (WB1: 20 mM Tris pH 7.5, 300 mM NaCl, 1 mM EDTA, 0.1% NP40) and transferred to a new tube. Beads were then washed 4 times with Wash Buffer 2 (WB2: 20 mM Tris pH 7.5, 300 mM LiCl, 1 mM EDTA, 0.5% NP40, 0.5% NaDOC) and transferred to a new tube. After a final wash with Wash Buffer 3 (WB3: 10 mM Tris pH 8, 300 mM NaCl, 1 mM EDTA), proteins were eluted with 200 μl of Elution Buffer (EB: 50 mM Tris pH 7.5, 300 mM NaCl, 10 mM EDTA, 1% SDS) at 65°C 30 min 1500 rpm in a ThermoMixer C (Eppendorf). The recovered eluate was then diluted with 1 volume of Elution Dilution Buffer (EDB: 50 mM Tris pH 8, 100 mM NaCl) and 0.1 mg of Proteinase K was added before crosslink reversal by incubating at 65°C for 10 hours. DNA was then recovered by phenol/chloroform extraction followed by ethanol precipitation in presence of 10 μg of glycogen. DNA ends were repaired and adenylated using NEBNext® Ultra™ End Repair/dA-Tailing Module according to manufacturer recommendations, followed by purification with ZYMO DNA clean & concentrator-5 kit. Illumina-compatible libraries were prepared using NEBNext® Multiplex Oligos for Illumina® (Index Primers Set 1) kit (NEB Cat# E7335S) according to manufacturer recommendations followed by purification with ZYMO DNA clean & concentrator-5 kit. DNA was eluted in 20 μl. Libraries were amplified in a 50 μl PCR reaction in the presence of 600 pmol of each Illumina universal forward primer and Illumina indexed reverse primers, using 5 μl of purified DNA as template and Phusion Hot Start DNA polymerase II (Thermo Fisher) according to manufacturer recommendations. After an initial denaturation step of 30s at 98°C, the PCR reactions underwent 16 PCR cycles (98°C 15s, 65°C 30s, 72°C 15s) and a final elongation step of 30s at 72°C. After purification of the PCR products on a ZYMO DNA clean and concentrator-5 column and elution in 10 μl, the PCR products were resolved on a 6% acrylamide/TBE gel. After staining the gel with SYBR Green 1, the area of interest was excised under blue light and PCR products were purified.

#### CLIP-Seq and CLAP-Seq

For the CLIP experiments, Dynabeads Protein G Magnetic Beads (1 mg) were washed two times for 1 min with Tris-buffered saline (TBS) containing 0.05% Tween-20 and coated with 5-10 μg of antigen-purified MERS1 antibody for 1 h in the same buffer. The beads were washed with PBS three times for 5 min. Antibody was crosslinked to IgG by adding 0.5 ml of freshly-prepared 0.45 mM DSS (disuccinimidyl suberate) solution in 1xPBS, and incubated for 1 h. The beads were pelleted and washed two times with 0.2 M glycine pH 2.5, and three times with TBS for 10 min.

For the CLAP experiments, the entire bottle of Rabbit IgG (100 mg, Sigma I5006-100MG) was resuspended in 7 ml of water and dialyzed against 2L of PBS overnight at 4°C. The recovered solution contained ~14 mg/ml of IgG. The entire contents of Dynabeads® M-270 Epoxy vial (300 mg) was transferred with 20 ml of 0.1M sodium phosphate buffer (pH 7.4) into 50 ml conical tube and incubated on Nutator for 10 min at room temperature. The suspension was divided equally between four 15 ml conical tubes and the beads were collected on a magnetic stand. IgG solution was centrifuged at 21,000g for 10 min, and coupling mixture was prepared by adding components in the following order: 3.5 ml of IgG were mixed with 9.85 ml of 0.1M sodium phosphate buffer; 6.65 ml of 3M ammonium sulfate was added gradually and mixed. After incubation for 5 min at room temperature, the mixture was filtered through 0.22 μm low protein binding filter. Coupling mixture (5 ml) was added to beads in each 15ml tube and incubated on Nutator for 20 hours at 30°C. Beads were collected on magnetic stand, rinsed with 10 ml of PBS three times by brief vortexing and quenched with 3 ml of 100 mM Glycine-HCl pH 2.5. The beads were collected and supernatant decanted as soon as possible. Beads were resuspended in each tube in 3 ml of 20 mM Tris-HCl, pH 8.8, collected into a single 15 ml conical tube and washed three times with 5 ml of 100 mM triethylamine pH 8.0 for 5 min at room temperature, 3 times with 10 ml of PBS for 5 minutes at room temperature, once with PBS plus 0.5% Triton X-100 for 5 minutes, then 3 times with 10 ml of PBS for 5 minutes. Beads were resuspended in 6 ml of PBS plus 0.02% sodium azide (50 mg/ml) and stored at 4°C for up to three months.

*T. brucei* cultures (1.6 L) were grown to ~20×10^6^ cells/ml. If TAP-tagged protein was expressed, cells were grown for ~72 hours post-induction. Cells were pelleted at 3000g for 15 min, washed with 50 ml of ice-cold PBS with 6 mM sucrose and resuspended in 32 ml ice-cold PBS with 6 mM sucrose. Half of suspension volume was distributed equally into 4 pre-chilled cover plates from 10 cm Petri dishes. The remaining half was kept on ice. Petri dishes were placed on 6 cm-tall cold blocks and irradiated three times at 400 mJ/cm^2^ in CX-2000 UV Crosslinker (UVP) with gentle mixing between UV cycles. Cells were transferred into 50 ml tube and 30 ml of PBS was used to collect the remaining material from Petri dishes. Crosslinked and control cells were collected at 3000g for 10 min and frozen with liquid nitrogen.

UV-irradiated and mock-treated cell pellets were resuspended in 3 ml of Extraction Buffer (EB, 50 mM Tris, pH 7.6, 150 mM NaCl, 5 mM MgCl_2_, 1% NP −40, 1/10 of Complete Protease Inhibitor tablet without EDTA, 40U of Turbo DNase) per gram of cells (wet weight) and incubated on ice for 15 min. Cells were diluted with 50 mM Tris, pH 7.6, 150 mM NaCl, 5 mM MgCl_2_ buffer to 11 ml and sonicated 3 times for 20 s at 12W with intermediate incubations on ice. The extracts were centrifuged at 40,000 rpm for 20 min in SW41 rotor. The supernatant was filtered through 0.22 μm low protein binding filter, EDTA was added to 10 mM, and the lysate was divided into two 15 ml conical tubes. Antibody-coated Protein G magnetic beads (1.5 mg) or rabbit IgG-coated Dynabeads (1 mg) were added per tube and incubated on Nutator for 30 min at 4°C. For High and Low RNase treatment, 2 and 0.2 μl of RNaseA/T1 cocktail (RNase A, 500 U/ml; RNase T1, 20,000 U/ml, Ambion) were added per tube, incubated at 26°C for 15 min, and placed on ice. After another incubation on Nutator for 30 min at 4°C, magnetic beads were rinsed with 10 ml of WB (20 mM Tris pH 7.6, 150 mM NaCl, 1 mM EDTA, 0.2% NP40) and washed two times with 10 ml of WB for 5 min. Beads were transferred into a 2 ml tubes, washed two times with 1 ml of WB in Thermomixer for 10 s, washed two times with 1 ml of HS buffer (20 mM Tris pH 7.6, 500 mM NaCl, 1 mM EDTA, 0.2% NP40) for 10 min.

For MERS1 CLIP and MERS2 CLAP, beads were washed two times with 1 ml of CIAP buffer (50 mM Tris pH 8.0) for 5 min, before adding 25 U of calf intestinal phosphatase in 50 μl of 1xCIAP buffer and incubating at 37°C for 15 min at 1000 rpm.

For MTRNAP CLAP, beads were washed with 1 ml of 50 mM Tris pH 7.5 buffer and incubated with 20 U of RNA 5′ polyphosphatase (Epicentre) in 50 μl of supplied buffer for 30 min at 37°C, 1000 rpm. After one wash with HS buffer and one wash with CIP buffer, beads were incubated with 20 U of RNase I (Epicentre) in a 50 μl reaction with the corresponding buffer for 30 min at 37°C for 30 min.

After one wash with WB, one wash with HS buffer, two washes with PNK buffer (40 mM Tris 7.5, 10 mM MgCl_2_) and one wash with 0.1 mg/ml BSA in water, the first RNA adapter ligation was set up in a 50 μl reaction containing 1xT4 RNA ligase buffer, 15 U of T4 RNA ligase, 1 mM ATP, 20 mg/ml BSA, 40 U of RNase OUT (Thermo Fisher) and 100 pmol of either RA3 3′ RNA adapter (MERS1 CLIP, MERS2 CLAP) or RA5 5′ RNA adapter (MTRNAP CLAP). Reactions were incubated overnight at 16°C, 1000 rpm.

After two washes with PNK buffer, RNA was radiolabeled with 10 μCi of [γ-^32^P] ATP, 10U of polynucleotide kinase in 50 μl of PNK Forward Buffer and incubated at 37°C for 10 min at 1000 rpm. Cold ATP was added to 0.2 mM, and beads were incubated for 20 more min. After two washes with 1 ml of WB and one with PNK, ribonucleoprotein complexes were eluted with 40 μl of 1xLDS-MOPS loading buffer with 50 mM DTT by incubating at 70°C for 10 min, 1000 rpm. After centrifugation of the supernatant for 5 min at 21,000g, eluates were resolved on a 4%-12% NuPAGE gel. Proteins were then transferred to nitrocellulose membrane in 1xMOPS buffer as recommended by manufacturer and exposed to phosphor storage screen. The membrane was stained with Sypro Ruby protein blot stain (Thermo Fisher) for 2-3 min and de-stained in water. The area of interest was then cut out under blue light into strips and incubated in 200 μl of PK buffer (100 mM Tris-HCl, pH 7.5, 50 mM NaCl, 10 mM EDTA) with 4 mg/ml of proteinase K at 37 °C for 20 min at 1000 rpm. The mixture was supplemented with 200 μl of 7M urea in 1x PK buffer and incubated for 20 more minutes. RNA was purified by adding 0.53 ml of phenol-chloroform (3:1, pH 5.2) and incubation for 20 min at 37 °C at 1000 rpm. After centrifugation at 21,000g for 5 min at room temperature, the upper aqueous phase was transferred to a new tube and RNA was precipitated with 50 μl of 3M Sodium Acetate (pH 5.2), 5 μg of glycogen and 1 ml of ethanol:isopropanol (1:1) mixture. After an overnight incubation at −20°C, RNA pellet was resuspended in water and the second RNA adapter was ligated in a 50 μl reaction in presence of 15 U of T4RNA ligase, 40 U of RNase OUT, and 20 pmol of the RNA adapter (RA5 for MERS1 CLIP and MERS2 CLAP, RA3 for MTRNAP CLAP) by incubating overnight at 16°C. After phenol/chloroform extraction and ethanol precipitation, cDNA was synthesized using SuperScript III with 3.5 pmol (MERS1 CLIP, MERS2 CLAP) or 20 pmol (MTRNAP CLAP) of RTP primer. Libraries were amplified in a 50 μl PCR reaction in the presence of 600 pmol of each Illumina universal forward primer and Illumina indexed reverse primers, using 5 μl of the cDNA synthesis reaction and Phusion Hot Start DNA polymerase II (Thermo Fisher). After an initial denaturation step of 30 s at 98°C, the PCR reactions underwent 16 PCR cycles (98°C 15 s, 60°C 30 s, 72°C 15 s) and a final elongation step of 30 s at 72°C. After purification of the PCR products on a ZYMO DNA clean and concentrator-5 column and elution in 10 μl, the PCR products were resolved on a 6% acrylamide/TBE gel. After staining the gel with SYBR Green 1, the area of interest was excised under blue light and PCR products were purified before sequencing.

#### In vitro activity assays

RNA substrates for *in vitro* enzymatic assays were prepared by *in vitro* transcription. The sequences were as follows:

ND8 mRNA fragment (maxicircle position 3165-3202):

GGAAUUUUUGGGGGAGCUCGACGGCGGGCGGAGCAUUAUUUGAGGAGGGCGGGA
GCAGAAGGCUUUCUGAGGAAAGAGGGGACCGAGAUCGAUGAAGGUUAUUUUUUG
GUUAUUGAGGAUUGUUUAAAAUUGAAUAAAAAGGCUUUUUGGAAGGGGAUUUUU
GGGGGACACCGCCAGAGGAGGAGGGUUUUGGAAGAGUUUGUUUU
Antisense RNA for ND7 mRNA (maxicircle position 4743-4674):

GGAGAAAGCCUUCUGCUCCCGCCCUCCUCAAAUAAUGCUCCGCCCGCCGUCGAGC
UCCCCCAAAAAUUCC

To generate DNA template for ND8 and anti-ND7 *in vitro* transcription, fully complementary primer pairs (C810/C811 for ND8 and C763/C764 for anti-ND7) were annealed by incubation at 95°C for 1 min and 37°C for 10 min with temperature ramping of −0.1 °C /s in Phusion DNA polymerase reaction mixture (Thermo Fisher) without polymerase and dNTPs. *In vitro* transcription templates of RPS12+ND5, ND7+CO3 and anti-RPS12 substrates were amplified from cloned maxicircle regions (positions 14276-16319 for RPS12+ND5 and anti-RPS12, and 3896-6944 for ND7+CO3) with Phusion DNA polymerase and primer pairs (C715/C756 for RPS12+ND5, C786/C758 for anti-RPS12, and C716/C762 for ND7+CO3). The amplification reaction was performed with 35 cycles at 95°C for 30 s, 55°C for 30 s, 68°C for 30 s. The duplex DNA templates were extracted with phenol-chloroform, precipitated in ethanol prior to *in vitro* transcription.

Triphosphorylated ND8 transcript bearing a 5′-terminal γ-^32^P was synthesized in 100 μl reaction containing 40 mM Tris-HCl pH 7.8, 20 mM NaCl, 26 mM MgCl_2_, 2 mM spermidine, 10 mM DTT, 0.1% Triton X-100, 80 U T7 RNA polymerase (Ambion), 1 μM template, 0.5 mM ATP, 0.5 mM CTP, 0.5 mM UTP, 12.5 μM unlabeled GTP and 0.42 μM [γ-^32^P] GTP (250 μCi, Perkin Elmer). After incubation at 37°C for 2 hours, the synthesized RNA was extracted with phenol/chloroform, precipitated with ethanol, and purified on 15% polyacrylamide/8M urea gel.

RPS12+ND5, anti-RPS12, ND7+CO3, and anti-ND7 transcripts were synthesized in a reaction containing 40 mM Tris-HCl pH 7.8, 20 mM NaCl, 26 mM MgCl_2_, 2 mM spermidine, 10 mM DTT, 0.1% Triton X-100, 0.8 U/μl T7 RNA polymerase, 1 μM template, and 4 mM NTP. After incubation at 40 °C for 3 hours, the synthesized RNA was extracted with phenol/chloroform, precipitated with ethanol, and purified on 10% polyacrylamide/8M urea gel electrophoresis. The purified transcripts (RPS12+ND5 and ND7+CO3) were dephosphorylated by recombinant Shrimp Alkaline Phosphatase (New England Biolab), and followed by 5′ labeling with T4 polynucleotide kinase (Ambion) in the presence of [γ-^32^P] ATP (Perkin Elmer). Labeled RNA was gel-purified.

Double-stranded RNA substrates were prepared by hybridizing 100,000 cpm of labeled RNA and 1 pmol of the complementary RNA in 10 μl mixture containing 50 mM Tris-HCl pH 8.0 and 100 mM KCl. RNA was annealed by incubation at 85°C for 2 min, 65°C for 5 min, and 20°C for 10 min with temperature ramping of −0.1°C /s.

Pyrophosphohydrolase activity assay was carried out in 20 μl reaction containing 50 mM HEPES pH 7.5, 100 mM NaCl, 2 mM MgCl_2_, 0.5 mM MnCl2, 1 mM DTT, 50,000 cpm of ^32^P-γ labeled ND8 substrate, and 4 μl of TAP-purified complexes fractions (MERS2, GRBC5, RGG2, and LSU-4710) or rapid affinity-purified MERS1 fraction. The reaction mixture was pre-incubated at 30°C for 10 min, and the reaction was started by addition of the RNA substrate. To identify the released pyrophosphate and orthophosphate, the same substrate was treated with 0.04 U/μl of 5′ RNA pyrophosphohydrolase (RppH, New England Biolab) in 20 μl reaction containing 10 mM Tris-HCl pH 7.9, 50 mM NaCl, 10 mM MgCl_2_, 1 mM DTT at 30 °C for 20 min. Aliquots (5 μl) were taken in 10, 20 and 30 min, and transferred into 2 μl of 2 μg/μl Proteinase K (Thermo Fisher) and kept on ice until last time point completed. All aliquots were incubated at 30°C for 10 min to digest proteins in the reaction, and 1.5 μl of each aliquot was analyzed by thin-layer chromatography on a PEI-cellulose F plate (Millipore) developed with 0.3 M potassium-phosphate buffer pH 7.4 for 1 hour. TLC plates were air-dried and exposed to phosphor storage screens. Phosphor images were acquired with Typhoon FLA 7000 (GE Healthcare).

*In vitro* MPsome processing activity. MPsome assay was carried out in 40 μl reaction containing 50 mM Tris-HCl pH 8.0, 1 mM DTT, 2 U/μl RNaseOut recombinant ribonuclease inhibitor (Life Technologies), 0.1mM MgCl_2_, 40,000 cpm of labeled ssRNA or dsRNA, and 4 μl from TAP-purified DSS1 or DSS1 DN fractions. The reaction mixture was pre-incubated at 30 °C for 10 min, and the reaction was started by adding RNA substrate. Aliquots (10 μl) were taken in 5, 10 and 20 min, transferred into 4 μl of 4×Native PAGE sample buffer (Life Technologies) supplemented with 1 μg/μl Proteinase K and kept on ice until last time point completed. All aliquots were incubated at 30 °C for 10 min to digest proteins in the reaction. To visualize assembled RNAs, 5 μl of samples was analyzed on 7% polyacrylamide/Tris-borate native gel (Acrylamide-bis Acrylamide 29:1). To detect only the labeled strand, 5 μl of samples were mixed with 5 μl of Stop Solution (95% formamide, 10 mM EDTA, 0.05% Xylene cyanol and 0.05% Bromophenol blue) supplemented with 100 nM of unlabeled substrate, heated at 85 °C for 2 min, incubated on ice for 5 min and separated on 10% polyacrylamide/8M urea denaturing gel (Acrylamide-bis Acrylamide 19:1). To measure the products lengths, the same RNA substrates were digested by guanine-specific RNase T1, and alkaline-hydrolyzed. RNase T1 reaction was carried out in 10 μl reaction containing 3 mM sodium citrate (pH 4.5), 0.1 mM EDTA, 0.8 M urea, 0.5 μg/μl yeast tRNA mixture (Ambion), 0.02 U/μl RNase T1 (Ambion), and 10,000 cpm of labeled ssRNA. The reaction was incubated at 55°C for 2 min 30 s and kept on ice. The equal volume of Stop Solution was added to the reaction prior to loading. Partial alkaline-hydrolysis was carried out in 10 μl reaction containing 100 mM sodium carbonate (pH 9.0) and 10,000 cpm of labeled ssRNA. The reaction was incubated at 95 °C for 2 min 30 s and kept on ice. The equal volume of Stop Solution was added to the reaction prior to loading. Gels were dried and exposed to storage phosphor screens. Phosphor images were acquired with Typhoon FLA 7000 (GE Healthcare).

### Quantification and statistical analysis

#### KAP-Seq

The raw pair-end sequencing data was and was in FastQ format. Adapter sequences (DNA and RNA oligonucleotide table) were trimmed from the raw reads by cutadapt (v1.14, https://pypi.python.org/pypi/cutadapt). Any read pair that contained <20 nt reads after adapter trimming was removed to suppress random alignment to reference genome. The read pairs were filtered against the *T. brucei* nuclear genome by using Bowite2 (v2.3.2, http://bowtie-bio.sourceforge.net/bowtie2/index.shtml) with default parameters. The reference genome was retrieved from TriTrypDB (Release 33, http://tritrypdb.org/tritrypdb/). The remaining read pairs were mapped to the maxicircle (Genbank ID: M94286.1) with default parameters. In-house Perl script #1 was used to calculate the actual length of the fragment using the SAM file generated by Bowtie2 as input. The script searched for reads pairs that concordantly mapped to maxicircle in forward orientation by interpreting the flag signal, and calculated the coordinates at which the reads were mapped on maxicircle by interpreting the CIGAR string. The length of an amplicon was defined as the distance between the start coordinates of read 1 and the end coordinates of read 2 in a concordantly read pair. The length distribution of the fragments was visualized as a histogram using in-house R-script. A similar in-house Perl script (#2) were used to read the SAM file from bowtie2 and calculate the read counts for every coordinate on maxicircle. The read count on a given coordinate was defined as the number of amplicons that overlapped with the coordinate. Read coverage on maxicircle was visualized using in-house R-script by plotting the maxicircle coordinates on the X-axis and the corresponding read counts on the Y-axis.

#### CLIP-Seq and CLAP-Seq

For the CLIP and CLAP experiments, we performed single-end 100 nt sequencing on HiSeq 2500 platform (Illumina). The stranded single-end sequencing data were supplied in FastQ format. Adapter trimming and filtering against nuclear genome were performed as above, and the remaining reads were mapped to the maxicircle genomic sequence and to edited mRNA sequences. In-house Perl script #2 was used to calculate the read counts mapped to each coordinate, for both the major strand and the minor strand of the maxicircle, and the read coverage on both strands was separately visualized with the in-house R-script. To test the correlation between MERS1 CLIP-Seq dataset and MERS2 CLAP-Seq dataset, we compared the CLIP/CLAP-Seq mapped read coverage per coordinate on both strands of the maxicircle and calculated a two-sided Pearson correlation score between the datasets. The total number of count pairs involved in the Pearson correlation test (N) was 46032. A correlation score p-value of less than 0.001 was considered significant.

The Motif analysis for MERS2 CLIP-Seq: The aligned BAM files were converted to BED format by bedtools bamtobed command (Quinlan and Hall, 2010). Then the peak calling was performed on major and minor strand separately by MACS 1.4.2 with the parameter setting “-keep-dup=all −shiftsize=1 −nomodel −g 24000” (Zhang et al., 2008). Motif calling was performed by MEME (Bailey and Elkan, 1994) only on the input strand, but not reverse complementary strand and using the maxicircle reference sequences as the background signal. The final motif was selected based on the distance to the peak location identified by MACS and the number of peak containing the predict motif.

#### RACE-Seq

For the 5′ RACE experiments, we performed 150 bp paired-end sequencing on the Illumina HiSeq 2500 platform. Procedures described above were used for adapter trimming, short read removal, nuclear contaminant removal and maxicircle read mapping. The SAM files generated by Bowtie2 was converted to sorted BAM files using SAMTools (v1.5, http://samtools.sourceforge.net/). The sorted BAM files were used to generate TDF files via the “count” command in igvtools (v2.3.95) with the following parameters: “-z 10 −w 1 --strand read” for temporary visualization on IGV (v2.3). Finally, by using the “tdf2bedgraph” command in igvtools, the TDF files were converted to BEDGRAPH files which contained per-nucleotide read counts like the output of Perl script #2. For the antisense 5′ RACE experiment, a modified version of the in-house R script was used to read BEDGRAPH files and visualize the read coverage for individual genes on maxicircle. In addition, the script also extracted the read counts from −100 nt to 20 nt of the 3′ end for every gene, and visualized the aggregation of the read counts for this region. For the sense 5′ RACE experiment, we had four biological replicates for each treatment group: parental, DSS1 dominant negative, mock, and polyphosphatase treated RNA. Pearson correlation were tested between each two samples among a total of 16 samples. We adopted the same method to calculate correlation scores as described for CLIP-Seq and CLAP-Seq. In all cases, the total number of count pairs (N) was 46032, and correlation scores with p-values less than 0.001 was considered significant. The correlation scores were summarized into a 16×16 table, and a heatmap visualization was generated by using the R-package corrplot (v0.77).

For the 3′ RACE experiment, we performed single-end 300 bp sequencing on the Illumina MiSeq platform. Procedures described above were used for adapter trimming, short read removal, nuclear contaminant removal. The remaining reads were mapped to the maxicircle using BWA (http://bio-bwa.sourceforge.net/). For reads with multiple alignments, we selected the alignment with the shortest 3’ unmapped region, which was defined as the tail of the transcript. The transcript tails were identified as “A-tails” or “U-tails” if the sequences were consisted of over 90% of the corresponding base. All other transcript tails were categorized as “other”. Reads without 3′ unaligned sequences were categorized as “No tail”. An in-house Python script was used to interpret the SAM file generated by BWA and calculate the read count and starting coordinate on maxicircle for each tail types defined above. The position of the tail for the ND9 gene was visualized with in-house R script.

**Appendix Table 1.**
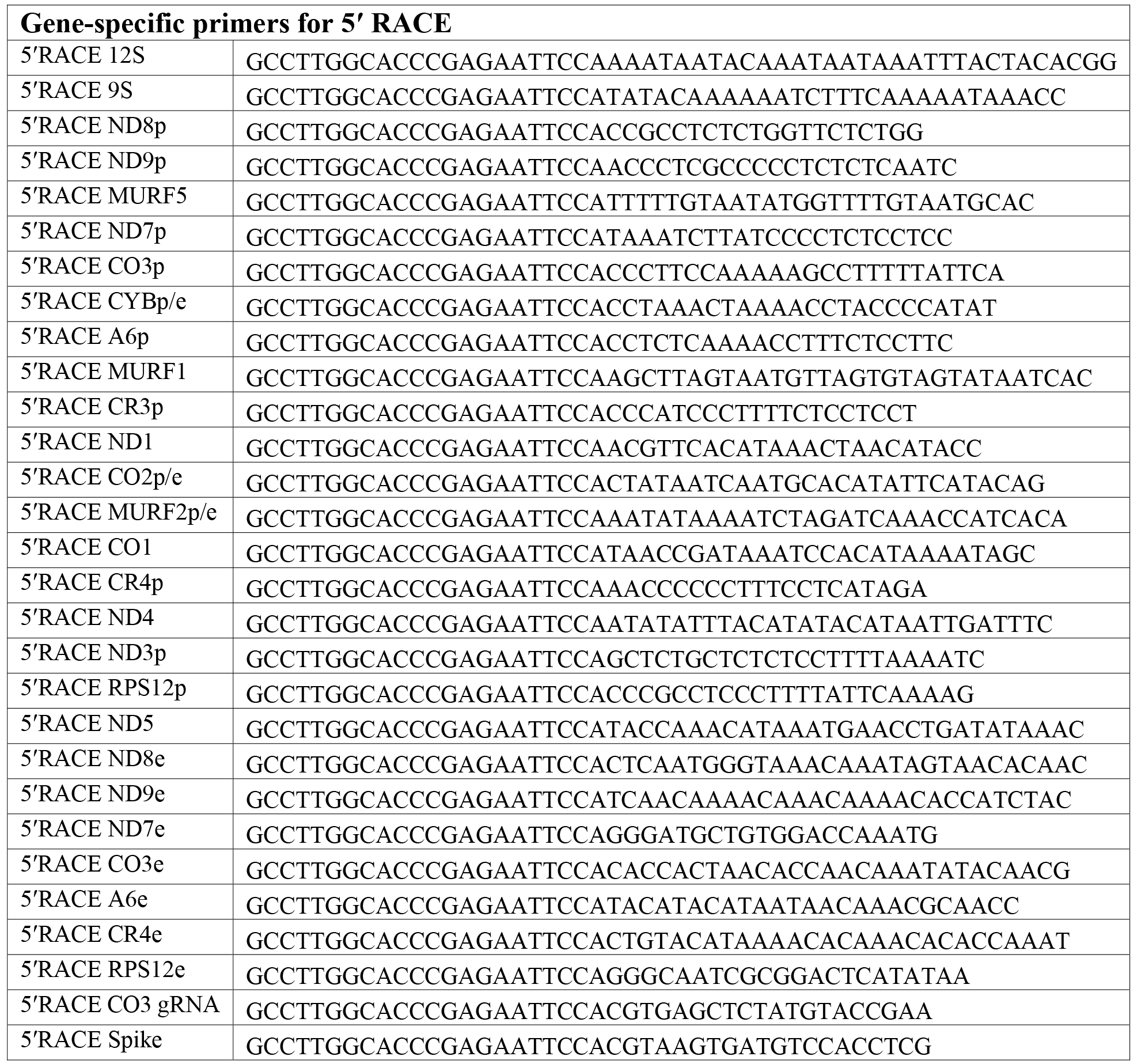
**DNA and RNA oligonucleotides.** p, pre-edited; e, edited; fw, forward; rv, reverse.

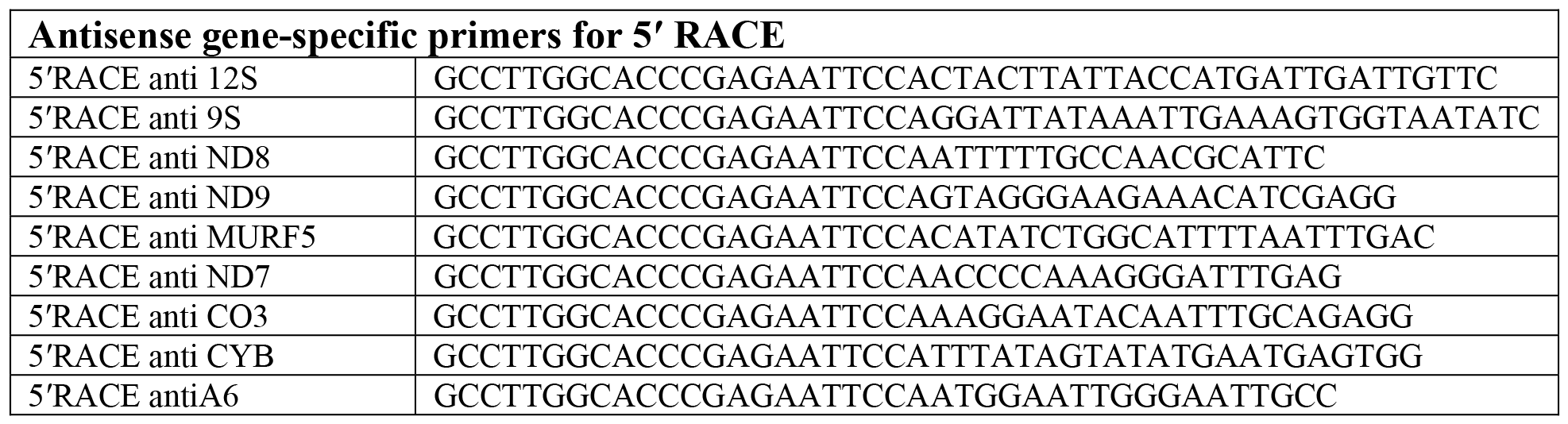

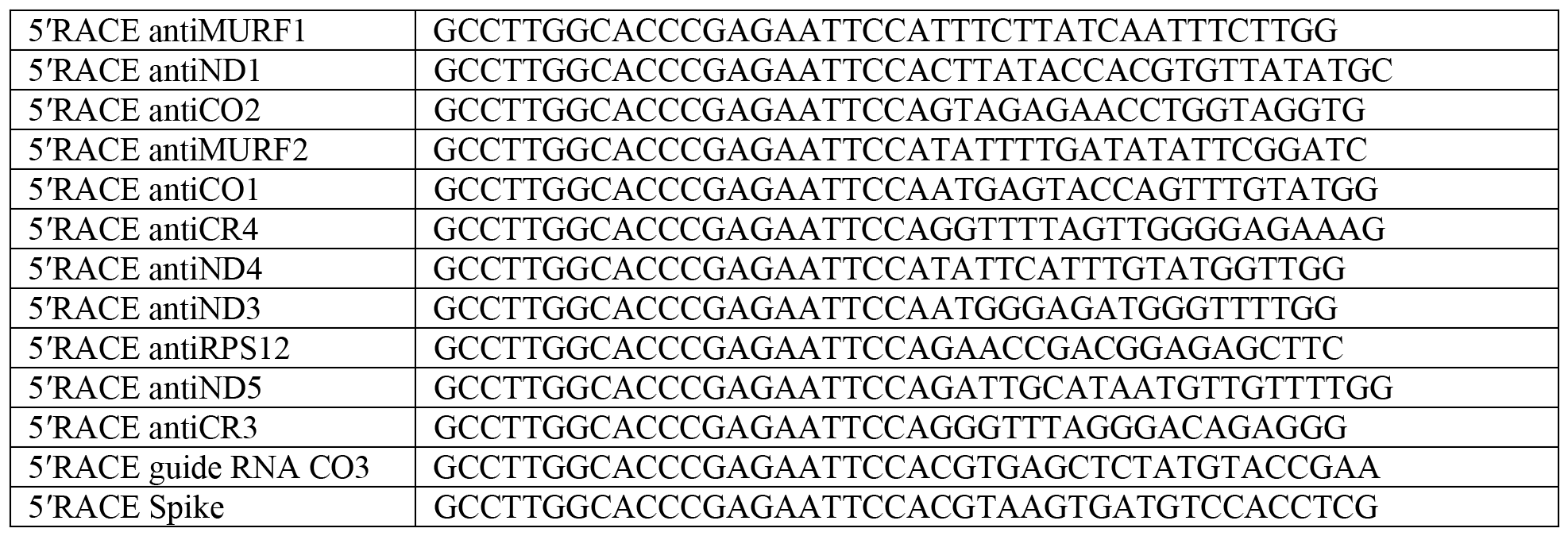

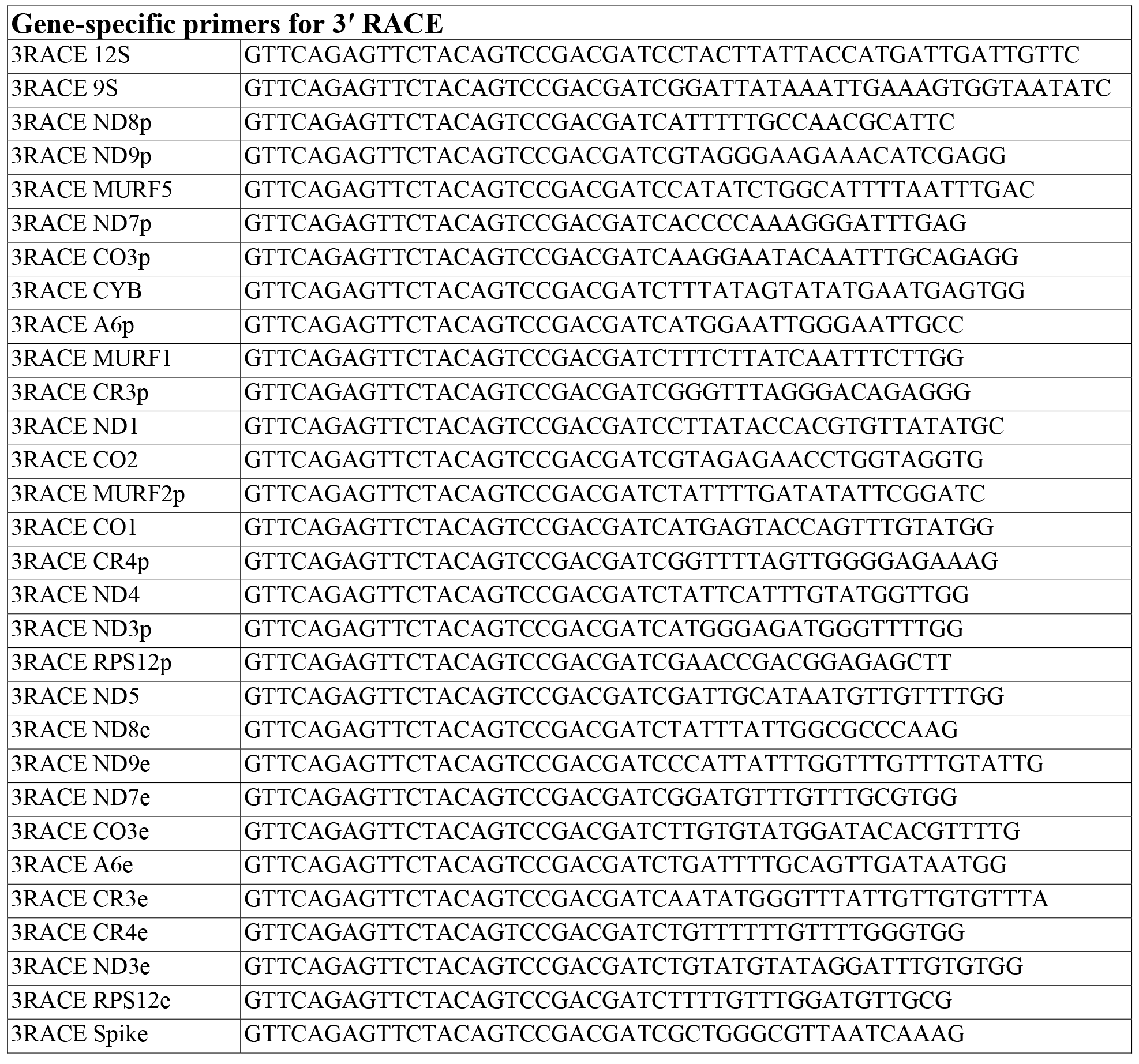

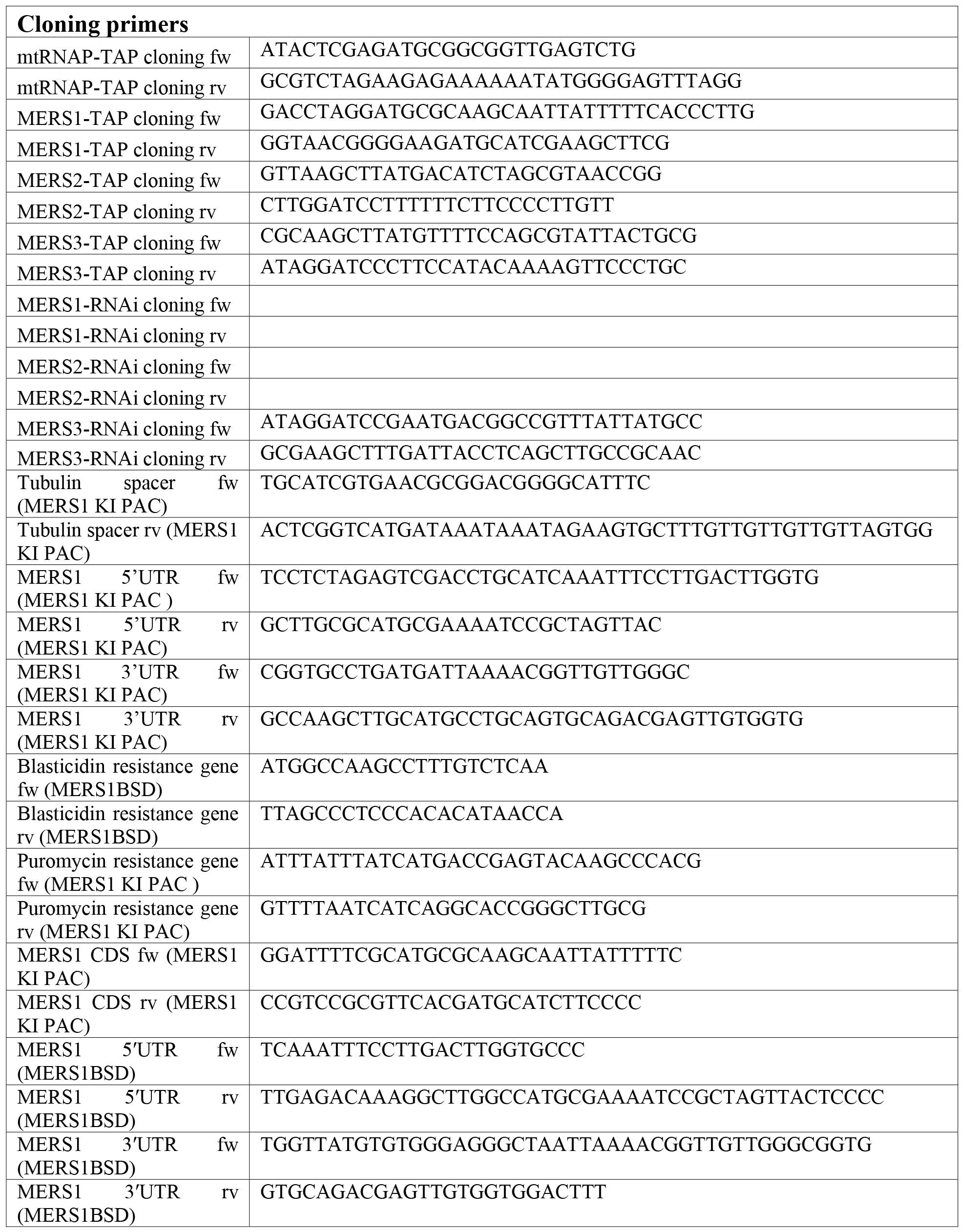

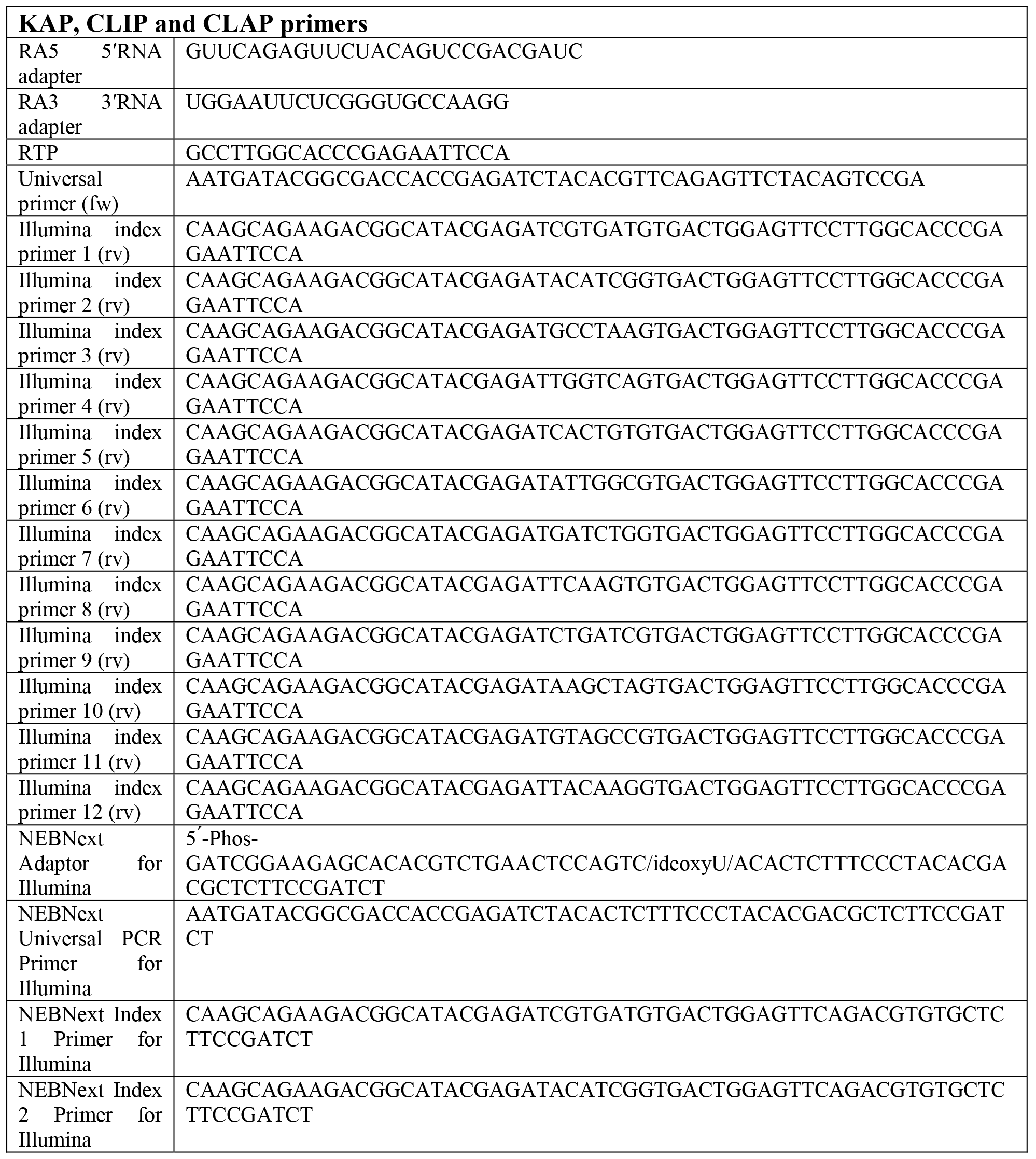

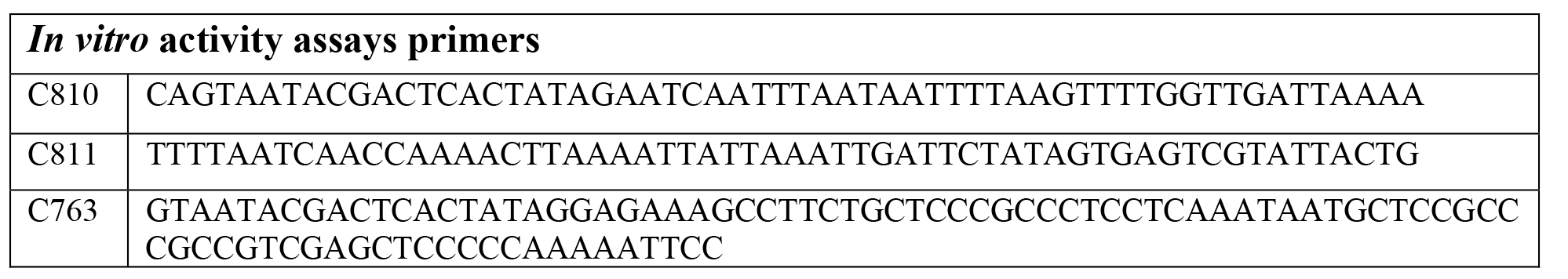

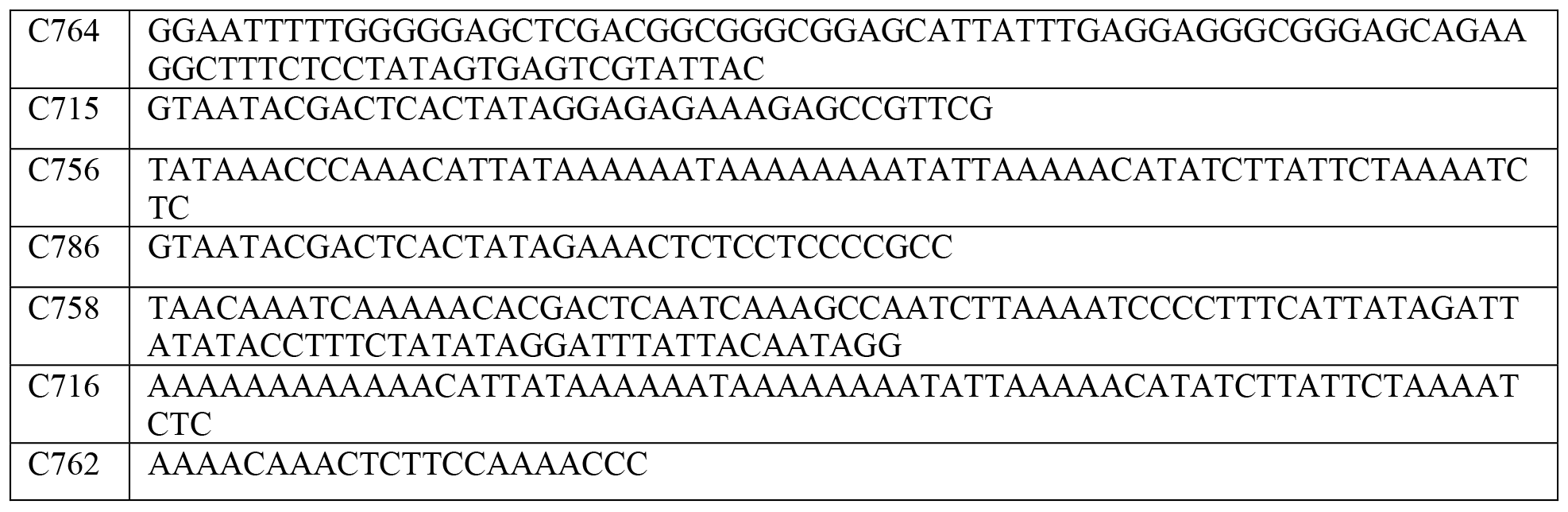

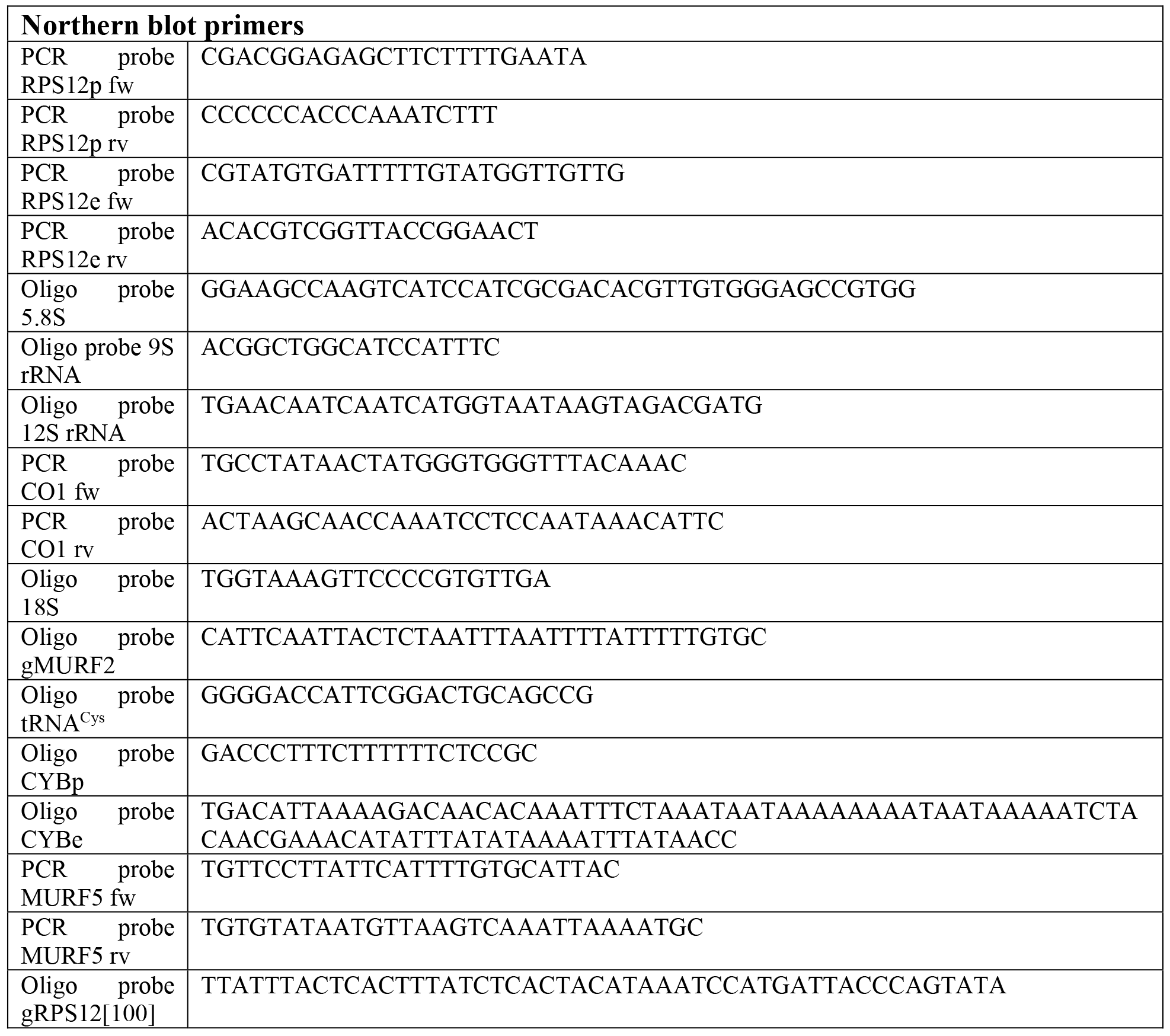

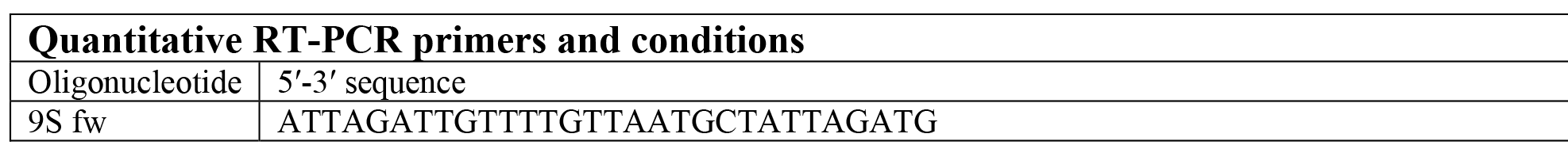

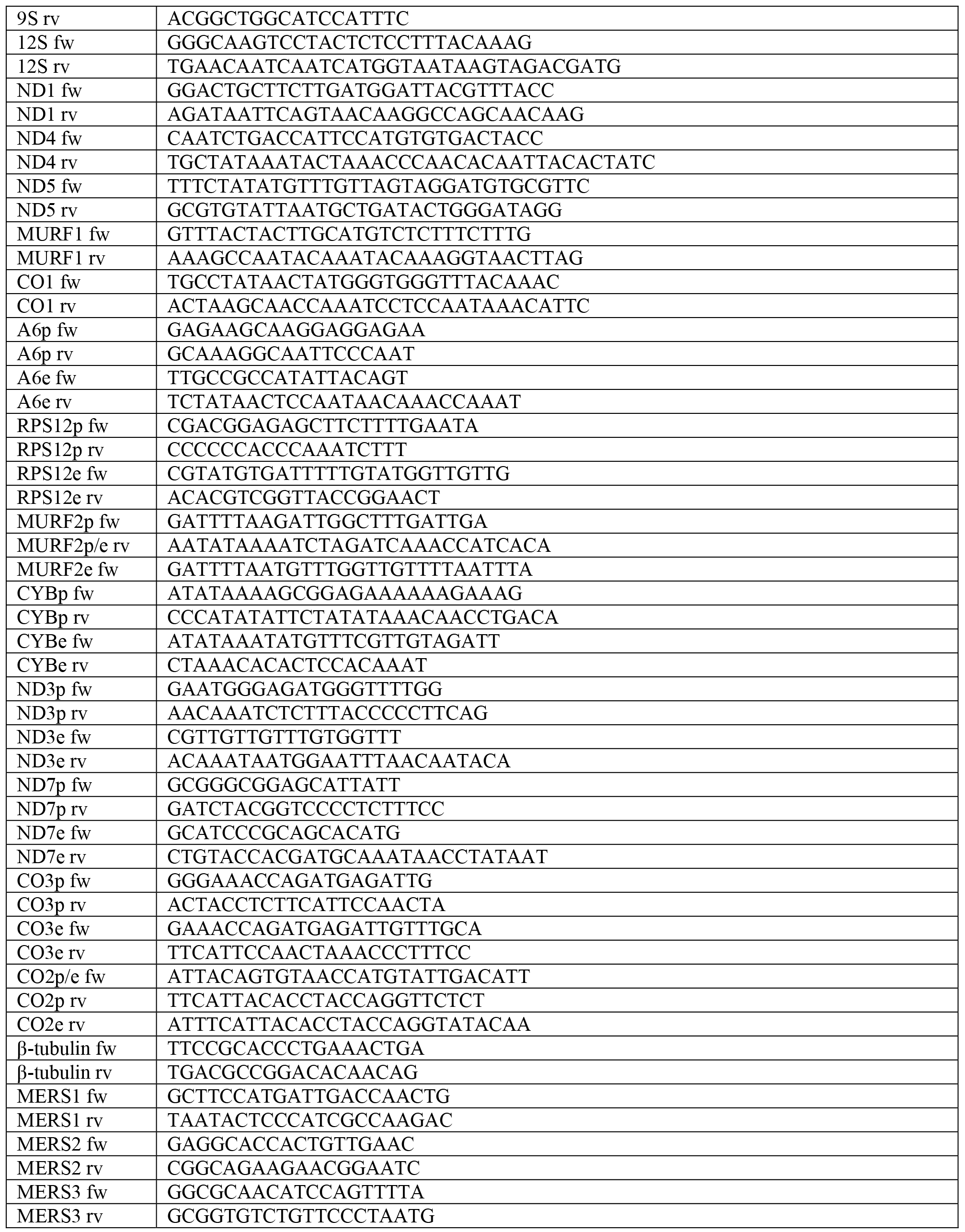

